# Persistent, Private and Mobile genes: a model for gene dynamics in evolving pangenomes

**DOI:** 10.1101/2024.07.15.603572

**Authors:** Jasmine Gamblin, Amaury Lambert, François Blanquart

## Abstract

The pangenome of a species is the set of all genes carried by at least one member of the species. In bacteria, pangenomes can be much larger than the set of genes carried by a single organism. Many questions remain unanswered regarding the evolutionary forces shaping the patterns of presence/absence of genes in pangenomes of a given species. We introduce a new model for bacterial pangenome evolution along a species phylogeny that explicitly describes the timing of appearance of each gene in the species and accounts for three generic types of gene evolutionary dynamics: persistent genes that are present in the ancestral genome, private genes that are specific to a given clade, and mobile genes that are imported once into the gene pool and then undergo frequent horizontal gene transfers. We call this model the Persistent-Private-Mobile (PPM) model. We develop an algorithm fitting the PPM model and apply it to a dataset of 902 *Salmonella enterica* genomes. We show that the best fitting model is able to reproduce the global pattern of some multivariate statistics like the gene frequency spectrum and the parsimony vs. frequency plot. Moreover, the gene classification induced by the PPM model allows us to study the position of accessory genes on the chromosome depending on their category, as well as the gene functions that are most present in each category. This work paves the way for a mechanistic understanding of pangenome evolution, and the PPM model developed here could be used for dynamics-aware gene classification.

## Introduction

Many bacterial species present an impressive diversity in terms of gene content. Due to pervasive intra- and inter-specific horizontal gene transfer (HGT), the number of genes present in a given species is often much higher than the number of genes contained in a typical genome of this species, and can potentially be very large. As multiple genomes from the same species began to be available around the year 2005, the observation that each new sequenced genome contained new genes lead Tettelin et al. (2005) to coin the term ‘pangenome’ for the set of all genes present in a species (Tettelin & Medini, 2020). Genes from a pangenome are usually classified as ‘core’ (if they are present in every genome) or ‘accessory’ (if they are not) (Page et al., 2015). The diversity and organization of bacterial genes could be used to understand the mechanisms governing their evolutionary dynamics. Understanding the dynamics of accessory genes is also of interest for public health as most genes conferring higher virulence or antimicrobial resistance belong to this category.

### Patterns and processes of pangenome evolution

In the genomic era, very rich information is available to study the patterns created by pangenome evolution: tens of thousands of sequenced genomes in certain bacterial species, including hundreds of completely assembled genomes. These patterns can be found in various summaries of the data. To name a few, the mechanisms of pangenome evolution probably influences the distribution of gene frequencies, the presence/absence patterns of genes, the diversity carried by gene families and the order of genes along a chromosome. For example, the gene frequency spectrum (GFS) - which is the histogram of gene frequencies in a set of genomes - often presents a ‘U’ shape that seems ubiquitous in bacterial pangenomes, whereby more genes are rare and very frequent than present at intermediate frequencies (Figure 5a). While these patterns can be precisely described and compared between different populations (Cummins et al., 2022; Botelho et al., 2023), an open problem is to identify the main processes generating them. To this end, we seek to develop a minimal model describing gene dynamics in a pangenome, that is able to reproduce the observed patterns. We started with three biologically relevant types of evolutionary gene dynamics, partially based on previous models of pangenome evolution, and tested several versions of our model to retain the one that was best able to reproduce these patterns.

### Existing models of bacterial pangenome evolution

Most models of bacterial gene evolution study gene gains and losses along a reference species tree, usually obtained using the alignment of core genome sequences. A simple model is the classical Markovian model of binary character evolution along a phylogenetic tree, in which a gene can be gained (0 → 1) and lost (1 → 0) at constant rates along the tree lineages (Pagel, 1997). Cohen and Pupko (2010) used this model to detect horizontally transferred gene families, by allowing the rates to vary across gene families and using stochastic mapping to map gain and loss events on branches. Using a similar model that allowed rate variation across time and lineages, and considering small genome chunks rather than genes as the evolutionary units, Didelot et al. (2008) inferred variations of genomic flux across lineages in several bacterial species and correlated it to their lifestyle. This model class was later called the Finitely Many Genes (FMG) model by Zamani-Dahaj et al. (2016). Indeed, this model assumes that a finite pool of genes is available to the population, which is fixed from the beginning of the species’ history, and remains constant throughout evolution.

Around 2012, alternative models emerged to account for gene immigration from other species. The Infinitely Many Genes (IMG) model (Baumdicker et al., 2012) assumes that new genes are imported at a constant rate along the phylogeny, then are lost at a constant rate and cannot be regained once lost. In this model, imported genes are picked from an infinite pool. Haegeman and Weitz (2012) presented a similar model with a fixed genome size, thus imposing that a gene importation must coincide with a gene loss. These models were originally designed to explain the U-shaped gene frequency spectrum (GFS) of pangenomes, and were fitted using exclusively the GFS.

Zamani-Dahaj et al. (2016) compared these two approaches and estimated the proportion of genes that were better explained by one gain (IMG model) versus multiple gains (FMG model), finding that both types of gene dynamics are present in 40 genomes of cyanobacteria. The two models are now implemented in the software Panaroo to allow for gain and loss rates estimation (Tonkin-Hill et al., 2020).

There are several limitations to the aforementioned approaches. First, some assumptions of existing models are limiting: the IMG model does not account for intra-species gene transfers, while the FMG model does not account for inter-species transfers. When comparing both types of gene dynamics, Zamani-Dahaj et al. (2016) find that under the FMG model, 15% of genes have patterns that are better explained by multiple gains, suggesting that they have been transferred. These genes are typically not well fitted by the IMG model. Meanwhile, the FMG model assumes that all genes from the pangenome are already present in the gene pool of the species from the root of the phylogeny. In reality, many genes are imported into the focal species over its evolutionary history, sometimes very recently compared to the whole evolutionary history. The IMG model has been extended to account for intra-species gene transfers and analysed to derive predictions on several pangenome summary statistics, but an inference method has not been developed yet (Baumdicker & Pfaffelhuber, 2014).

We also identify methodological limitations. On the one hand, studies aiming at fitting the GFS use a very restricted fraction of the available data (the gene frequencies). As a result, they are not able to reproduce correctly the level of parsimony observed in gene presence/absence patterns. On the other hand, while Zamani-Dahaj et al. (2016) reproduce the GFS of 40 cyanobacteria species thanks to several types of gene dynamics and allowing variability in parameters, the best model has five categories of genes. With much larger genomic datasets available, it is necessary to define parsimonious models that lend themselves more easily to interpretation. In this direction, we suggest a middle ground approach. We fix a priori the number of categories and parameters based on biological observations of gene dynamics, and check how well we are able to reproduce some of the global patterns observed in the data to decide whether more complex models are needed.

### Persistent, Private and Mobile gene dynamics

In the following, we list three qualitatively distinct gene behaviours that are relevant for bacterial populations. Our goal is to design a model of pangenome evolution accounting for these three types of gene dynamics.

First, we model the dynamics of genes that are in most circumstances essential to the bacteria, and thus are present in the ancestor and in the majority of sequenced genomes (Figure 1a). We use the adjective ‘persistent’ for these genes, as we expect them to largely overlap the set of genes usually designated as persistent (i.e., genes being present in a high percentage of genomes, typically between 95 and 100%: Perrin and Rocha, 2021). We hypothesise that the inferred loss rates of these genes will be very small. A loss can happen for example when a taxon colonizes a new niche where this gene is no longer essential, or by gene redundancy following the acquisition of a new gene performing the same function.

**Figure 1:**
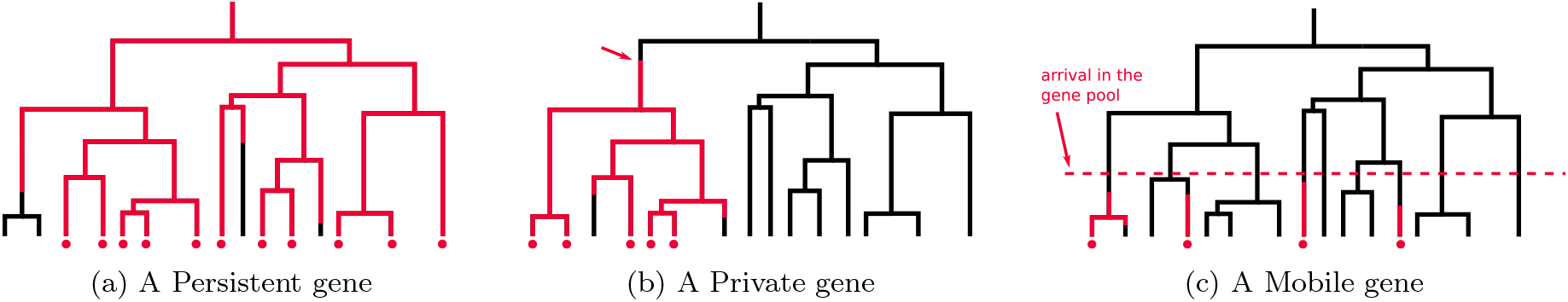
The three behaviours captured by our model: (a) Persistent genes are present at the root of the phylogeny and rarely lost (b) Private genes arrive once in the phylogeny, transmit vertically and are occasionally lost, while (c) Mobile genes arrive once in the gene pool of the population, then undergo intra-species HGT (thus multiple gains and losses along the phylogeny).

Second, we model genes that are specific to a given clade (Figure 1b). This specificity can have several explanations: a single event of transfer into the focal species, adaptation to a particular environment (e.g. bovine-adapted *Escherichia coli* lineages: Arimizu et al., 2019), defence against a phage specific to this clade (e.g. in *Vibrio crassostreae*: Piel et al., 2021), or epistatic interactions with genes already present in this clade (Whelan et al., 2021; McInerney, 2022; Beavan et al., 2024). We call such genes ‘private’. Some of these genes are already present in the ancestor of the population, while others appear along the species tree either by *de novo* gene birth or by inter-species transfer.

Finally, we model the dynamics of ‘mobile’ genes, i.e. genes that undergo frequent intra-species HGT such as transposons, prophages or genes located on plasmids. These genes have highly non-monophyletic presence/absence patterns at the leaves of the species tree (Figure 1c). A mobile gene is first introduced into the gene pool of the population by inter-species transfer, which is a rare event, and then is able to spread in the population through frequent intra-species HGT (Tenaillon et al., 2010). For example, the *bla*_CTX-M_ gene family encodes CTX-M *β*-lactamases allowing their host to tolerate *β*-lactam antibiotics. Originally found as a chromosomal gene in genus *Kluyvera*, it has introgressed several times in other Enterobacteriaceae species and has rapidly spread within these species on various mobile genetic elements (D’Andrea et al., 2013).

In this paper we introduce the PPM model for pangenome evolution. It includes three generic types of evolutionary gene dynamics along a species phylogeny, designed to model the aforementioned behaviours: (i) Persistent genes are present in the ancestral genome and can only be lost, (ii) Private genes are either present in the ancestral genome or introduced once in an internal lineage, are lost sometimes and undergo no further transfers, and (iii) Mobile genes undergo intra-species transfers following their arrival in the gene pool. We inferred the parameters of our model on a dataset of 902*S. enterica* genomes by fitting the presence/absence patterns of genes. Using these parameter values, we simulated genomes evolving according to our model and compared the obtained patterns to the ones from observed data. By assigning each gene to its most likely category, we were able to study the distribution of the three categories along the chromosome and on plasmids, as well as the gene functions present in each category.

### The model

In the Persistent-Private-Mobile (PPM) model, genes evolve on the species phylogeny along which they can be gained and lost. The following events are accounted for: gene gain and loss, intra- and inter-specific horizontal gene transfer, sequencing and bioinformatics errors. In particular, gene duplication and the simultaneous transfer of multiple genes are not accounted for in our model. These limitations are further discussed at the end of this paper. For the sake of concision, we index the three gene categories of the model as follows: 0 for the Persistent category, 1 for the Private category and 2 for the Mobile category. Parameters referring to each of these categories are indexed accordingly.

- *N*_0_ Persistent genes are present in the ancestral genome. They are lost at constant rate *l*_0_ along the phylogeny.
- *N*_1_ Private genes are present in the ancestral genome. At constant rate *i*_1_, new Private genes arrive on branches of the phylogeny. Once they are present, they are lost at constant rate *l*_1_ along the phylogeny. In the following, we fixed *N*_1_ to be equal to the expected number of Private genes at stationarity, which is *i*_1_*/l*_1_ (Huson & Steel, 2004). This category follows exactly the IMG model (Baumdicker et al., 2012).
- Mobile genes enter the gene pool of the population at constant rate *i*_2_ across time (i.e. along the tree height). Once they are in the gene pool, Mobile genes can be gained and lost at constant rates *g*_2_ and *l*_2_ by all lineages of the population phylogeny below their arrival date. Gains are interpreted as transfers from the pool to a tree lineage. The fact that we only model this type of transfers (rather than transfers between lineages) is consistent with the predominance of so-called ‘transfers from the dead’: assuming that the number of sequenced genomes is small compared to the total number of individuals in the species (which is usually the case), a transferred gene has much more chance to come from a non-sampled (‘dead’) lineage than from another lineage of the reconstructed tree (Szöllosi et al., 2013). One limit to this approach is that the gain rate is constant and identical for all Mobile genes, it does not depend on the frequency of each gene in the population.

These events are represented in Figure 2. Note that the PPM model requires a time-calibrated tree, because it defines at what points in the tree a mobile gene can be gained, given its arrival time. It thus assumes that gain and loss rates are approximately constant in time, as they happen at a constant rate on a dated tree.

**Figure 2:**
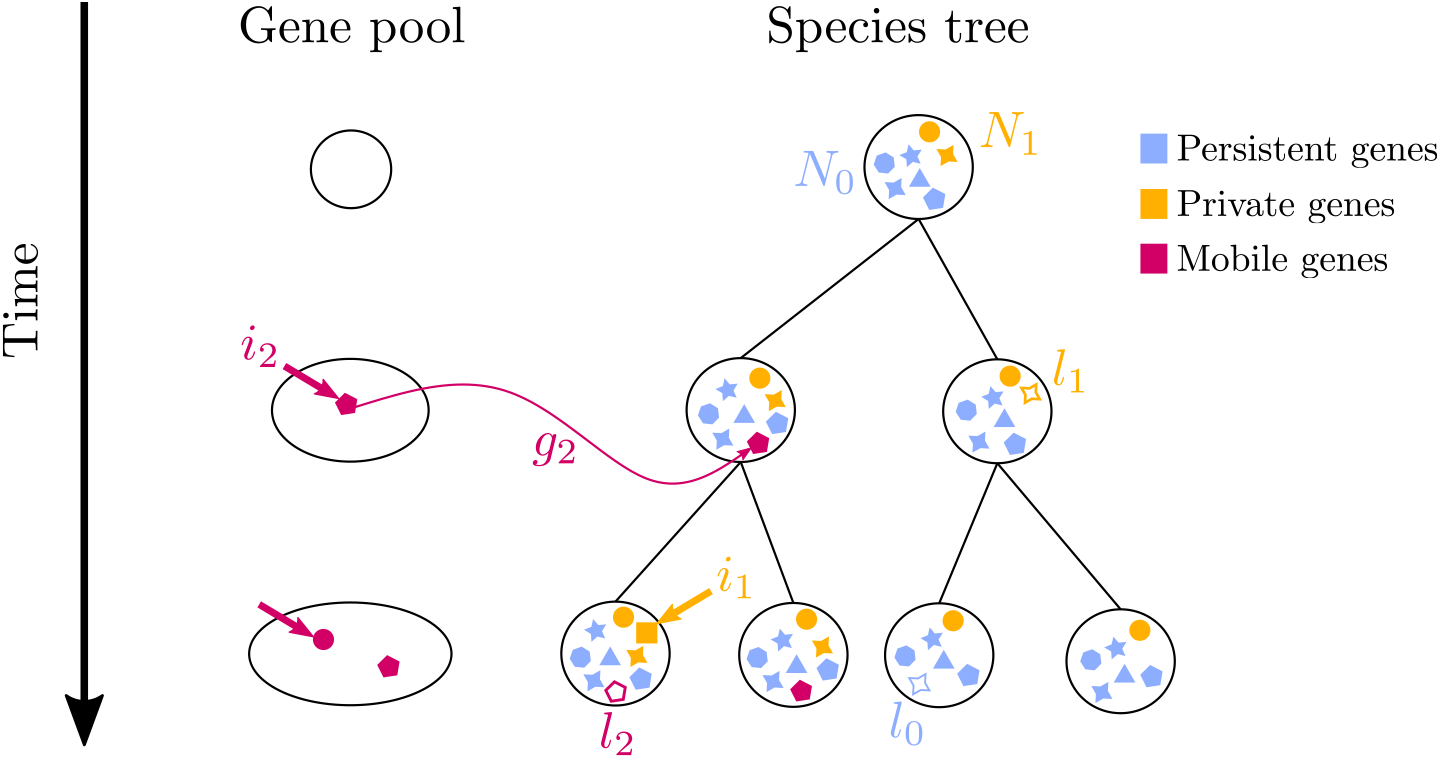
Schematic description of genomes evolving according to the PPM model for pangenome evolution. Persistent genes are represented in blue, Private genes in orange and Mobile genes in Burgundy. Different shapes represent different gene functions. The following events are represented, for Persistent genes: loss (rate *l*_0_); for Private genes: immigration on the tree (rate *i*_1_), loss (rate *l*_1_); and for Mobile genes: immigration into the gene pool (rate *i*_2_), gain (rate *g*_2_) and loss (rate *l*_2_).

Lastly, our model accounts for instantaneous switching between gene presence and absence at the leaves of the phylogeny. We added this possibility in the model for three main reasons. First, pangenome datasets can be subject to several sources of errors (Tonkin-Hill et al., 2020). These sources include sequencing errors (Salzberg, 2019) and bioinformatic errors (Denton et al., 2014). Second, genuine biological process of rapid gain and loss events of genes on the leaves of the phylogeny could be accounted for by this ‘switching’ term. For example if a bacterium having recently acquired a very deleterious gene is sequenced, it would probably not have left descent over several generations of replication *in vivo* (which is the case of bacteria at internal nodes of the phylogeny). This illustrates that gain and loss events may happen at the leaves at an evolutionary scale which is different from that of the rest of the phylogeny. Last, instantaneous switching at the leaves generates less parsimonious gene patterns and improves the fit to the data (Figure S8). Different parametrizations of switching probabilities are discussed in section 4 of the Supplementary Text. We present in the main text the results of inference for the model with two switching parameters *s*_1→0_ and *s*_0→1_. The parameter *s*_1→0_ is the probability to switch from presence to absence at a leaf for all gene categories, assumed to represent sequencing and bioinformatics errors. We found that we had little signal in our data to estimate a rate of rapid loss at the leaves for each category (Supplementary Text 4). The parameter *s*_0→1_ is the probability to switch from absence to presence, for Mobile genes only (“rapid gain”).

Our model does not explicitly include selection acting on gene presence/absence, although the interpretation of the dynamics of gene categories can involve selection: Persistent genes are under purifying selection, Private genes can interact epistatically with the genetic background of the clade, Mobile genes can be favourable in some environmental conditions but not in others.

The three gene categories that we model here have qualitatively distinct behaviours, except for Persistent genes and Private genes present at the root, who share the same behaviour with just different loss rates. Statistical justification for these three categories (rather than two) and exploration of rate heterogeneity inside categories is discussed in section 5 of the Supplementary Text.

## Results

### Simulation study

In order to assess the accuracy of our inference procedure (see Methods section), we simulated sets of presence/absence patterns and checked if we were able to infer the parameters that generated them. We randomly picked a 200-leaf tree, and 100 sets of values for parameters *N*_0_, *l*_0_, *i*_1_, *l*_1_, *i*_2_, *g*_2_, *l*_2_, *s*_1→0_ and *s*_0→1_. For each set of values we simulated genes evolving along the tree, and used the resulting presence/absence patterns to compute the maximum likelihood parameters. More details on this simulation study are given in section 2 of the Supplementary Text. Figure 3 shows the relations between the true parameters (i.e. those used for simulations) and the inferred ones.

**Figure 3:**
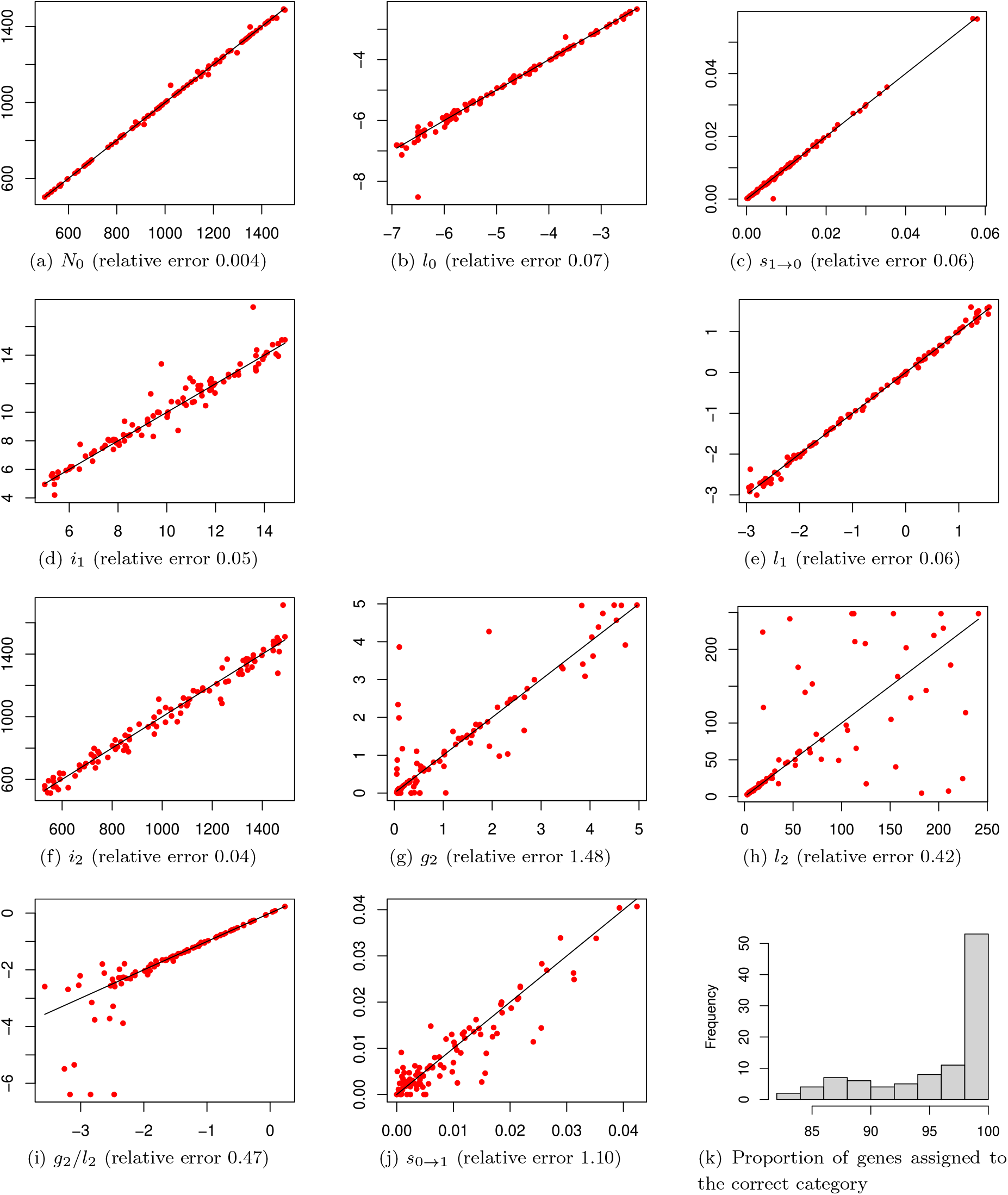
Results of the cross-validation study on a 200-leaf tree. (a-j) show correlations between true values (*x*-axis) and point estimates (*y*-axis) of parameters used to simulate sets of presence/absence patterns. Correlations for gain and loss parameters (*l*_0_, *l*_1_, *g*_2_*/l*_2_, *g*_2_ and *l*_2_) are plotted on a log scale. The legends of subplots display the mean relative error for each parameter. The mean relative error is computed by taking the absolute value of the difference between true and inferred values, divided by the true value, and then averaging over the 100 parameter sets for each parameter. (k) is a histogram showing the proportion of patterns that were assigned to the correct category by our method across the 100 simulated sets.

Most parameters show good correlations, with relative errors below 10%. In particular this is true for parameters governing the number of genes in each category: *N*_0_, *i*_1_ and *i*_2_. The switching parameter *s*_0→1_ shows higher relative error (110%). The gain and loss rates for the Mobile category are not very well estimated with relative errors of 1.48 and 0.42. This is due to very poor inference when *l*_2_ is more than 100 times bigger than *g*_2_ (Figure 3i). We discuss this phenomenon in section 2.3 of the Supplementary Text, and show on Figure S4 that this ratio should be well estimated in the case of the *S. enterica* dataset. The average proportion of genes correctly assigned to their category is 96% across the 100 simulated sets (Figure 3k).

### Maximum likelihood estimates of parameters

We inferred the Maximum Likelihood parameters of our model on a dataset of 902 *S. enterica* genomes, and found the following estimates:

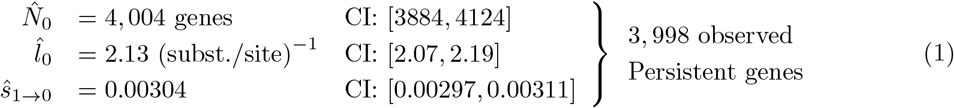

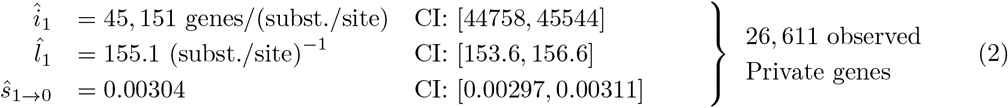

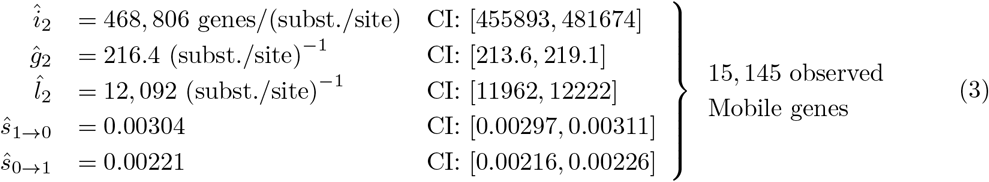

CI are asymptotic 95%-confidence intervals, computed using the second derivatives of the likelihood at its maximum. From these estimates, we computed several quantities of interest. The predicted number of genes in the ancestral genome is 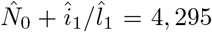 (CI: [4169, 4421]), close to the mean number of genes per genome in the dataset (4,556).

The expected number of genes of each category in the observed pangenome is computed using the forthcoming set of equations (6), and we find an expected number of 3,998 Persistent genes, 26,611 Private genes and 15,145 Mobile genes. This means that the observed pangenome is composed of 9% of Persistent genes, 58% of Private genes and 33% of Mobile genes, as shown on Figure 4. We also assigned each gene to its most likely category, and computed the proportion of each category present in an average chromosome and in an average plasmid (Figure 4). As Persistent genes are typically present in many genomes, they represent a small proportion of the observed pangenome (9%) but a high proportion of the genes carried by a chromosome (83%). The opposite holds for Private and Mobile genes: they compose a small proportion of the chromosome (11% and 6%, respectively) but the majority of the pangenome. A typical genome from our dataset carries 4,556 genes, of which 3,699 are Persistent, 491 are Private and 365 are Mobile.

**Figure 4:**
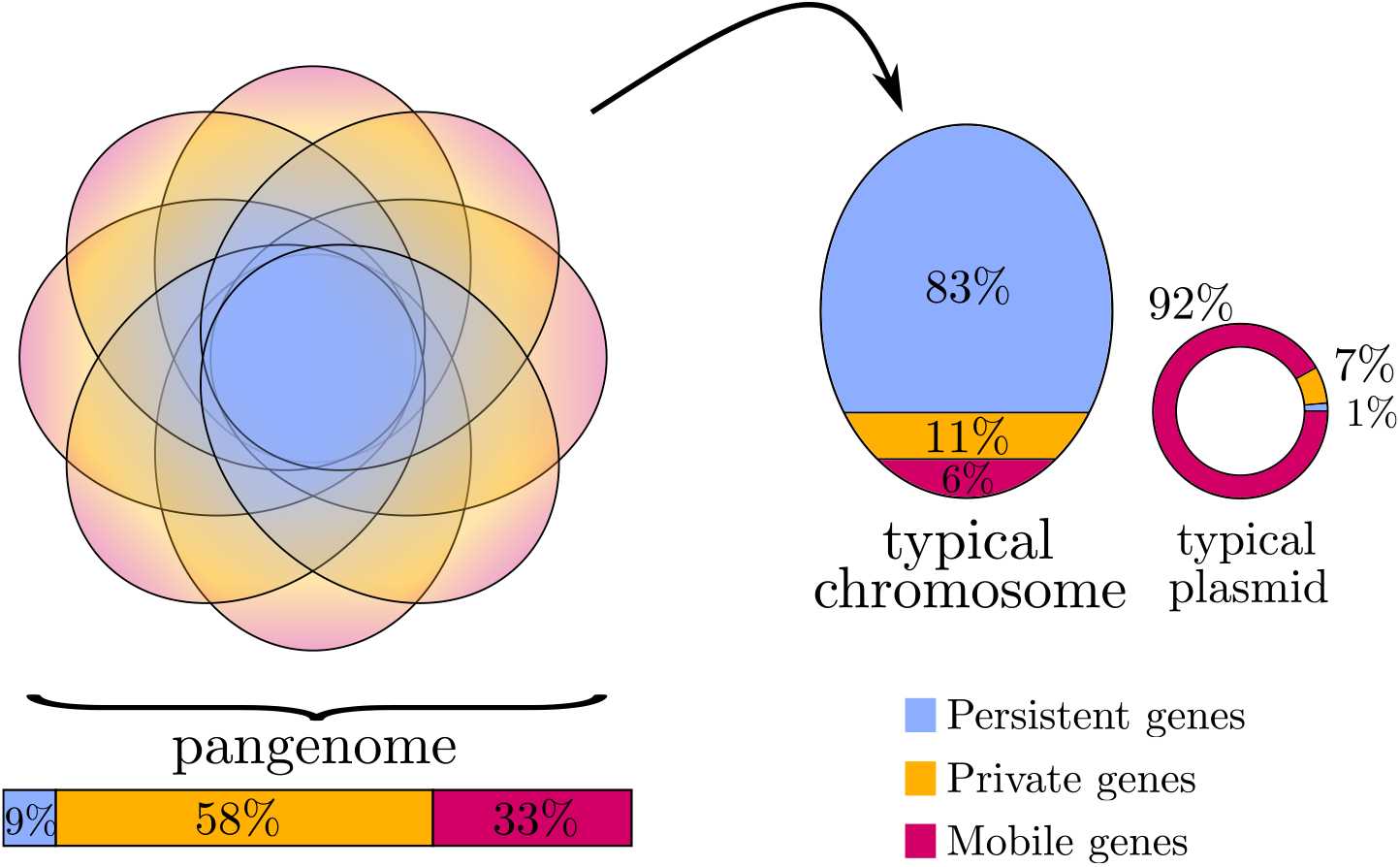
Proportion of the Persistent, Private and Mobile categories in our *S. enterica* dataset as inferred by our model. Colours indicate gene categories: blue for Persistent, orange for Private and Burgundy for Mobile. The observed pangenome is composed of 9% of Persistent genes, 58% of Private genes and 33% of Mobile genes. A typical chromosome carries 83% of Persistent genes, 11% of Private genes and 6% of Mobile genes, while a typical plasmid carries 1% of Persistent genes, 7% of Private genes and 92% of Mobile genes. The Venn diagram on the left is for illustration purposes and does not represent real data.

**Figure 5:**
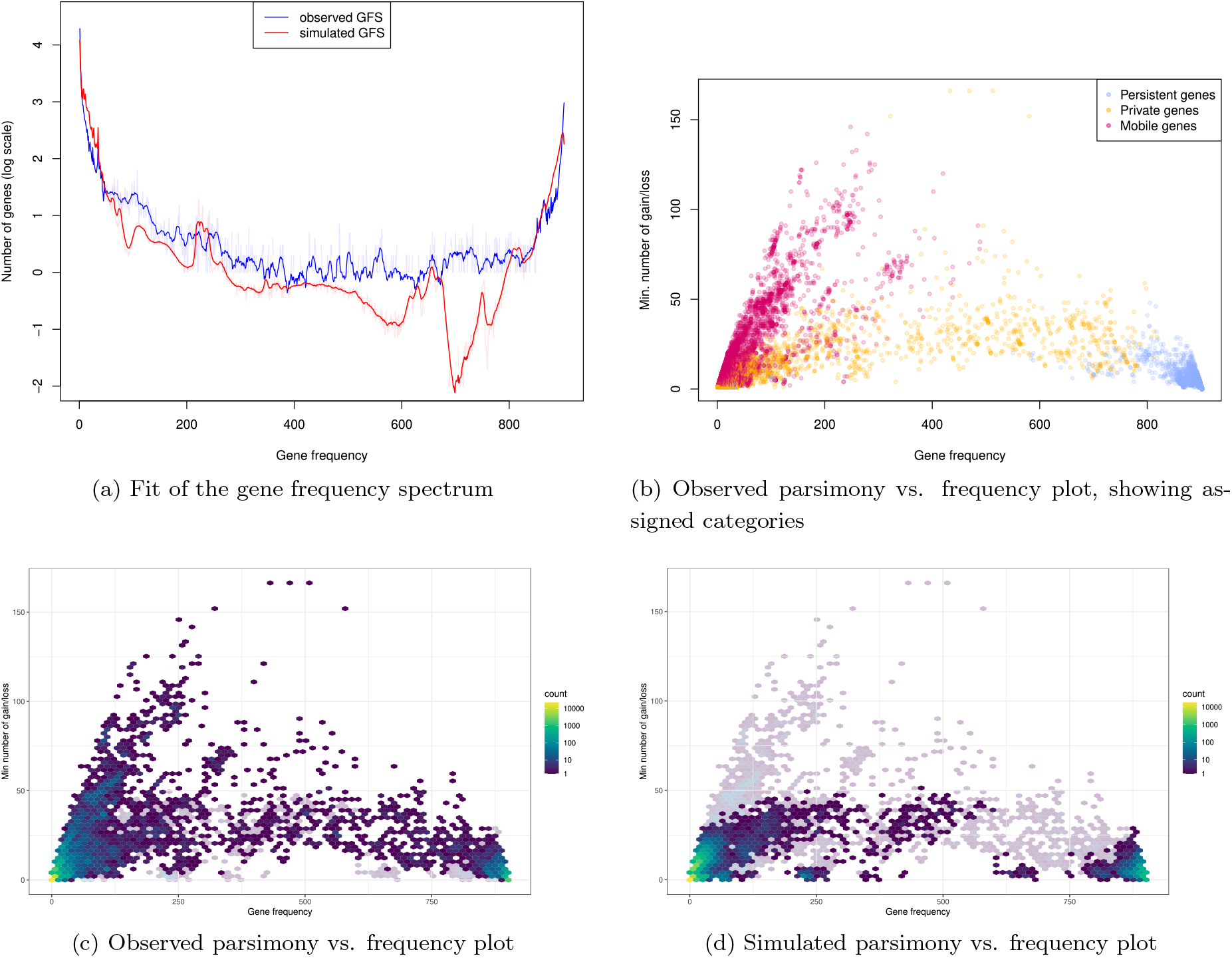
Two multivariate summary statistics of pangenomic data that we use in order to assess the goodness-of-fit of our model. Observed data are 902 genomes of *S. enterica*. Simulated data were obtained by simulating a set of genes evolving according to our model with Maximum Likelihood parameters estimated on observed data. (a) shows the histogram of gene frequencies, called the Gene Frequency Spectrum (GFS). Observed data are plotted in blue, while simulated data are in red. The simulated GFS is an average over 100 simulations. Curves are smoothed for better visualization; raw curves are shown in transparency. (b), (c) and (d) show the parsimony vs. frequency plot: the *y*-axis is the minimum number of gain and loss events needed along the species tree to explain the presence/absence pattern of a gene with frequency given on *x*-axis. (b) shows the observed parsimony vs. frequency plot coloured according to assigned categories. (c) shows the observed parsimony vs. frequency plot coloured according to points density, with simulated data in transparency. Conversely, (d) shows the simulated parsimony vs. frequency plot with observed data in transparency.

The loss rates are ordered as we expected, increasing from Persistent to Private to Mobile gene categories (although from the simulation study we know that the absolute value of *l*_2_ may not be very well inferred). Their values can be compared with the average branch length of the tree, which is 5 *×* 10^−4^ substitutions per site – keeping in mind that the tree has been transformed to be ultrametric, thus assuming a strict molecular clock. This length corresponds to a few thousand years if we divide by the estimated mutation rate of 10^−7^ substitutions per site per year for *S. enterica* (Duchêne et al., 2016). During this time interval, Persistent, Private and Mobile genes have a probability 0.001, 0.07 and 0.98 to be lost respectively. Additionally, a Mobile gene has a probability 0.02 to be gained if it was absent. These probabilities were computed using the transition matrix shown in equation (15).

During these thousand years, a given genome is expected to gain 23 Private genes that may result from *de novo* gene birth and direct transfers from other species, and 761 genes from the pool (if we assume that we are in the middle of the tree and the pool has around *i*_2_ *× H/*2 genes, where *H* is the tree height). The large probability of loss and large number of new gains of Mobile genes over thousand years implies that Mobile genes completely turnover over that period of time. By construction, a Private gene is introduced only once in the phylogeny. On the other hand, Mobile genes can be introduced several times from the gene pool to lineages of the phylogeny. We computed the expected number of distinct introductions for a Mobile gene using forthcoming equation (23), and found that they are introduced 26 times on average.

The switching probabilities are quite small (between 0.2 and 0.3%). This may indicate that the presence/absence patterns at the leaves are primarily driven by the underlying processes and not by random switching at the leaves.

### Goodness of fit on two summary statistics: the GFS and parsimony plot

While we fitted our model using the presence/absence matrix, we verified that it is also able to reproduce two important patterns of summary statistics characterizing the observed data. To that end, we simulated a pangenome using our model with the maximum likelihood parameter values inferred on the *S. enterica* dataset, and compared patterns from observed data versus simulated data.

The first pattern is the gene frequency spectrum (GFS), i.e. the distribution of gene frequencies in the observed pangenome. We chose the GFS as a summary statistic because it was used to fit pangenome evolution models in several previous studies. We plotted on Figure 5a the observed GFS (in blue) as well as the simulated GFS (in red). While the two curves do not perfectly correspond, in particular with a deficit of medium-high frequency genes in our model, the model qualitatively reproduces the U-shape found in the data.

The second pattern is the parsimony vs. frequency plot, which shows the parsimony of gene patterns as a function of their frequency. The *y*-axis is the minimal number of gains and losses needed along the species tree to explain a gene’s presence/absence pattern, thus small values correspond to very parsimonious patterns. We represented on Figure 5c the observed parsimony plot and on Figure 5d the simulated one. Our model reproduces the qualitative patterns of this plot, but the presence/absence patterns it generates are more parsimonious than in the observed data. In particular, the maximum score reached by simulated genes is around 50 while some observed genes go as high as 150. For a more detailed inspection of this parsimony vs. frequency plot, we plotted separate plots for each category on Supplementary Figure S7.

### Inferred categories

Using the ML parameter estimates, we computed for each gene the category with highest posterior probability (see equation (22) in Methods). We assigned each gene to a category – Persistent, Private or Mobile. We then compared different characteristics of the three categories, such as frequency, parsimony, position on the genome and enrichment in certain gene functions. The ternary plot on Supplementary Figure S3 shows that the assigned categories are clear-cut, with genes clustered around the corners of the triangle.

Figure 5b shows the parsimony plot coloured according to assigned categories. Genes assigned to the Persistent category have typically very high frequency, i.e. they are positioned on the right of this plot.

Surprisingly, we also spot a few genes at low frequency that are assigned to the Persistent category. Taking a closer look at their presence/absence patterns revealed that these genes are present at high frequency in two or more small clades. As a consequence, they are not well-fitted by the Private or the Mobile category. Genes assigned to the Private category present a wide range of frequencies, from very low (the majority of singletons is ‘Private’) to intermediate and high frequencies. While we could have expected singletons to be assigned preferentially to the Mobile category, it is not the case as Mobile genes have a tendency to be introduced several times (and thus, to be present in several genomes). Genes assigned to the Mobile category have low to intermediate frequencies and represent the majority of genes having low parsimony (i.e., in the upper part of the parsimony plot).

### Gene position

We studied the spatial distribution of the three gene categories on the bacterial chromosome and on plasmids. About two thirds of the studied *S. enterica* genomes contain one or several plasmids. On Figure 4 is represented the relative proportion of the three categories on a typical chromosome versus on a typical plasmid from the *S. enterica* dataset. As expected, we observe that chromosomes carry a majority of Persistent genes (83%), with some Private and Mobile genes (11% and 6%, respectively). On the other hand, plasmids carry almost exclusively Mobile genes (92%), which is consistent with their easy transfer.

We looked more closely at the spatial distribution of genes along a chromosome, starting arbitrarily with the first genome of the dataset, represented on Figure 6a. Gene are coloured according to their assigned category (Persistent in blue, Private in orange, Mobile in Burgundy). In accordance with the relative category proportions described above, we observe a majority of Persistent genes on the chromosome. Moreover, Private and Mobile genes are not uniformly distributed but appear clustered.

**Figure 6:**
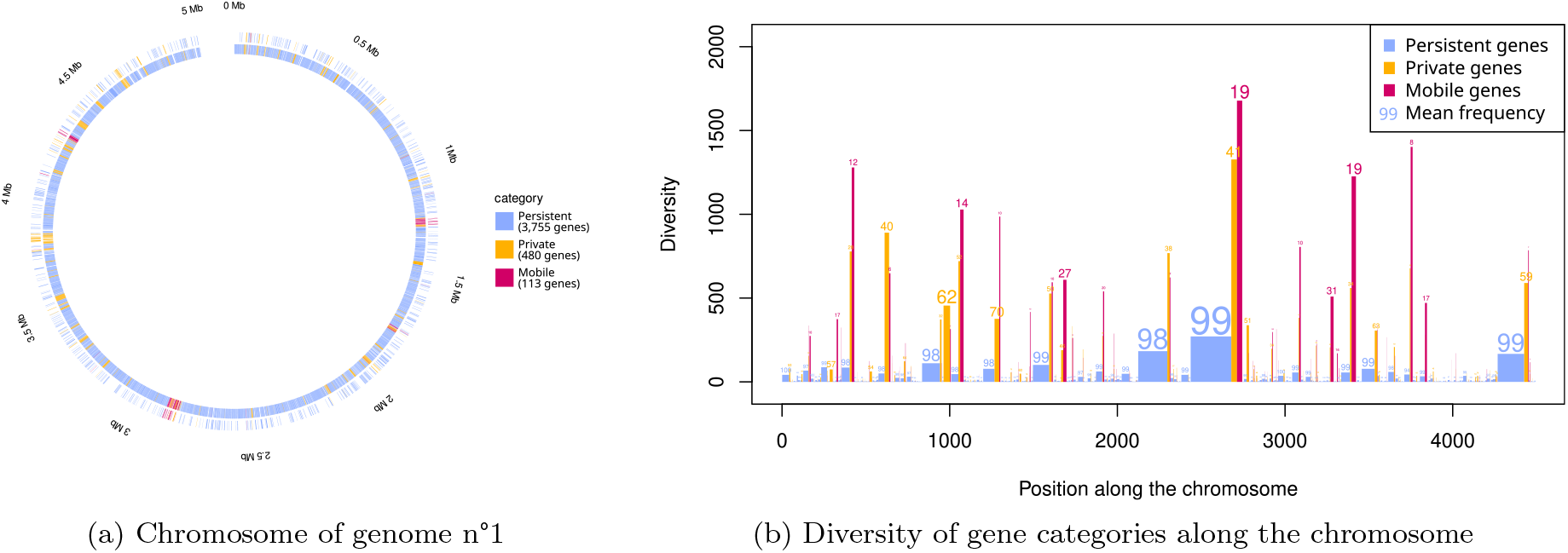
Distribution of gene categories along the chromosome. (a) shows gene categories along the chromosome of the first genome (chosen arbitrarily) of the *S. enterica* dataset. (b) shows the diversity of gene categories along the *S. enterica* chromosome, computed over 641 genomes sharing the same ordering of core genes. The position along the chromosome refers to the order of genes. Each bar represents genes from a given category present in a given interval between core genes. The width of the bar is the mean number of genes of this category present in this interval, and the height is the number of unique genes of this category present in this interval (i.e. the diversity). The number above the bar is the mean frequency of these genes.

To investigate whether the structure in gene category along the chromosome and the hotspots of Private and Mobile genes are conserved across strains, we used the 770 core genes as reference positions. We identified 641 genomes from the *S. enterica* dataset that had the exact same ordering of core genes. For each genome and each interval between two core genes, we computed the number of genes of each category that were present in this interval. We illustrate the diversity and distribution of gene categories along the chromosome of these 641 genomes with conserved synteny (Figure 6b). Hotspots of Private and Mobile genes that are conserved across the genomes appear all along the chromosome. These hotspots include from one gene to a few dozen genes with low frequency, but across all genomes carry a substantial gene diversity. Notably, the six most diverse hotspots of Mobile genes each carry more than 1000 unique genes across the 641 genomes. Mean diversity of Private genes in an interval is strongly correlated with mean diversity of Mobile genes in the same interval (Pearson correlation coefficient 0.91). The correlation is weaker for the mean number of genes of each of these categories in a given interval (0.58). The concentration of accessory genes in a few hotspots is in agreement with a study from Oliveira et al. (2017), where the authors find that genes imported by HGT are concentrated in hotspots representing around 1% of the genome in a dataset comprising 80 bacterial species.

### Gene functions

In order to study the gene functions present in each category, we selected genes that were annotated with a Cluster of Orthologous Genes (COG) database function (Galperin et al., 2021). A total of 2,743 Persistent genes, 5,612 Private genes and 2,639 Mobile genes were annotated with a function, representing respectively 70%, 21% and 18% of these categories. From this we built an association table between assigned gene category and COG function, illustrated on Figure 7. Overall, there is a highly significant association between gene category and gene function (*p <<* 10^−6^).

**Figure 7:**
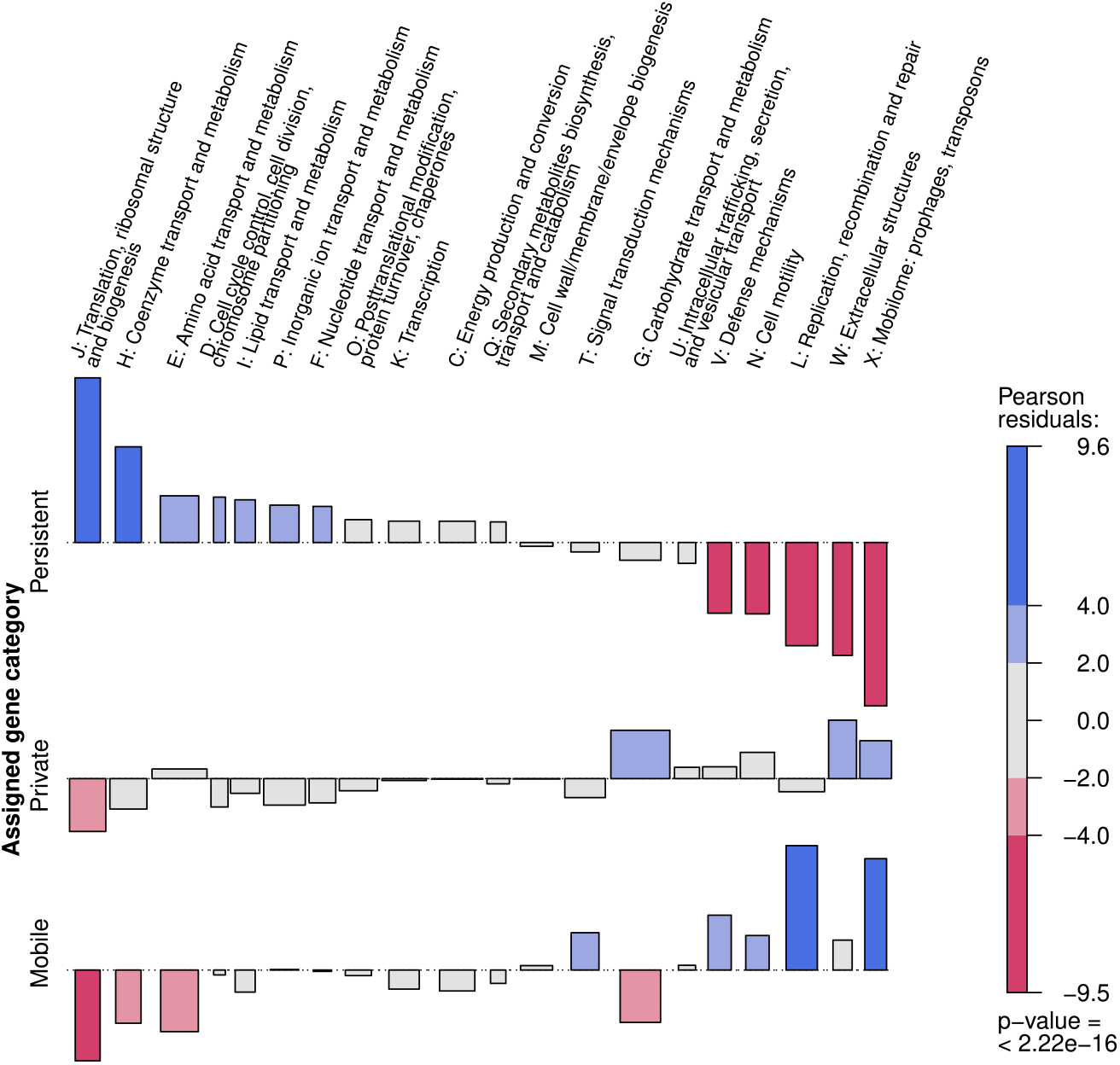
Association between inferred gene categories (Persistent, Private, Mobile) and COG functions. Rectangle heights represent the Pearson residuals, i.e. the observed number of genes corresponding to a given pair of category and function, minus the expected number of such genes if category and function were independent, divided by the square root of the expected value. Rectangle widths are proportional to the square root of expected values, so that rectangles areas are proportional to the difference between observed and expected value. A positive residual indicates a positive association between a gene category and a gene function, while a negative one indicates the opposite.

In particular, there is a positive association between the Persistent category and key cellular functions such as translation (J), cell cycle control (D) and the transport and metabolism of coenzymes (H), amino acids (E), lipids (I), inorganic ions (P) and nucleotides (F). On the contrary, the Private and Mobile categories have nearly opposite associations to the Persistent genes. They are both positively associated with the mobilome (X). The Private category is also associated with carbohydrate transport and metabolism (G) and extracellular structures (W). In addition, the Mobile category is positively associated with replication, recombination and repair (L), defence mechanisms (V), signal transduction mechanisms (T) and cell motility (N). The fact that many genes involved in ‘replication, recombination and repair’ are inferred to be Mobile genes is probably due to transposases (enzymes helping moving transposons) annotated as recombinases.

## Discussion

In this paper, we introduce the “Persistent, Private, Mobile” model for bacterial pangenome dynamics. This model proposes to classify bacterial genes into three qualitatively different dynamics on a phylogenetic tree, accounts for both intra- and inter-species gene transfer, and is scalable to large datasets. The model fitted to a dataset of 902 *S. enterica* genomes reproduces the U-shaped gene frequency spectrum and the parsimony plot, a new representation of gene configuration on a phylogeny. Moreover, the category assignment is clear-cut and the behaviours that we intended to capture in the three gene classes are consistent with the inferred gain and loss parameters, as well as with the function and position of genes in the genome. This model could therefore be used for dynamics-aware gene classification.

### A model for dynamics-aware category assignment

The PPM model was intended to model three gene categories with different behaviour and characteristics. The Persistent category was designed for housekeeping genes present at high frequency. Indeed, we find that genes assigned to this category have a low loss rate and a high mean frequency (853). They are inferred to form the majority of genes present on the chromosome of *S. enterica*. Moreover, this category is enriched in essential functions such as translation, cell cycle control and metabolism. The Private category was designed for accessory genes that are specific to a given clade. Genes assigned to this category have an intermediate loss rate and a small mean frequency (16). Interestingly, they are inferred to form the majority of the observed pangenome, mainly because singletons are classified as ‘Private’ and therefore represent 71% of this category. The ‘Private’ category is enriched in genes linked to carbohydrate transport and metabolism, and to extracellular structures. These two associations could be explained by a lasting clade-specific adaptation to different nutrients and environments. Studies performed on *S. enterica* have shown that different serovars display specific carbohydrate metabolism pathways (Seif et al., 2018) and adhesive mechanisms (e.g. pili and adhesins, see Wagner and Hensel, 2011). This category is also enriched in genes from the mobilome (transposons, prophages), which may be driven by singletons. The Mobile category was designed for accessory genes undergoing frequent intra-species HGT. Genes assigned to this category have a high loss rate and a low mean frequency (22). They represent the overwhelming majority of genes found on plasmids. This category is positively associated with gene functions that are known to be highly transferred: prophages, transposons, transposases, defence mechanisms. It is also associated with signal transduction mechanisms and cell motility.

Overall, our classification identifies persistent genes, genes adapted to certain clades and highly mobile genes. This classification cannot be obtained solely based on the gene frequency, even when additionally considering parsimony. For example, Private and Mobile genes can have similar frequency and parsimony (Figure 5b), but are actually organized differently on the species phylogeny. The strength of our classification of bacterial genes is that it is informed by patterns and relies on processes. The fact that it uses the gene’s inferred dynamics to classify it allows for a straightforward interpretation. Comparatively, the classification done by the tool PPanGGOLiN also uses the presence/absence matrix, but relies only implicitly on processes by assuming that neighbouring genes have more chance to share the same category (Gautreau et al., 2020).

Classifying genes on the basis of their dynamics may provide clues as to how genes of interest spread. This has also enabled us to study the distribution of different types of genes along the chromosome and on plasmids. As a result, we have identified hotspots of accessory gene insertion (Private and Mobile) along the chromosome of *S. enterica* (Figures 6a and 6b). In addition, dynamics-informed classification is potentially of great interest in comparative pangenomic analysis, providing an easy-to-interpret overview of the types of dynamics at work in different populations or species.

### Limitations

Our model has several limitations. First, it does not account for gene duplications. Although the method we use to generate gene families allows them to have multiple members per genome, we ignore duplicated genes by taking a binarized version of the presence/absence matrix. Thus, duplicated genes are treated as one gene even if they are sometimes present in multiple copies. Families containing at least one duplicate represent 8% of all families, and those containing at least two duplicates are only 3% of all families. We verified with additional simulations that gene duplication happening at rates compatible with our data does not alter parameter inference (Supplementary Text 6).

Second, we do not model the simultaneous transfer of multiples genes. We expect that sets of genes transferred in chunks influence our likelihood more than they should because they are counted as resulting from independent evolutionary events, while they are in reality co-transferred. However, this effect could be limited (Kloub et al., 2021). Kloub et al. designed a method called HoMer to detect potential horizontal multi-gene transfers (HMGTs). In a dataset of 103 *Aeromonas* genomes, they detected that 8.5% of potential intra-species gene transfers were contained in an HMGT event (this percentage was much higher for inter-species transfers, of which 20% were estimated to be part of a HMGT). This does not necessarily mean that HMGTs are rare, but rather that most genes involved in these transfers were removed by drift or natural selection before present and thus are not detected. Figure S9 shows a heatmap of pattern dissimilarity among accessory genes along a chromosome from the *S. enterica* dataset. Light squares on the diagonal reveal several groups of genes that could have been transferred together, as they are inserted in the same spot and show similar presence/absence patterns across the whole dataset. This visualization could help define groups of co-transferred genes. Our inference algorithm could then be re-ran on only one representative gene per chunk. However, as the composition of chunks differ across genomes, the problem of defining chunks across the whole dataset remains to be addressed.

Last, the importance of gene transfers in bacteria rises the question of how reliable are phylogenetic reconstructions, and what they represent. Studies have shown that a bacterial species tree can be reliably inferred even in the presence of gene transfers and conversions (in *Escherichia coli* : Touchon et al., 2009), and that phylogenetic distance is well correlated with gene content (in *Acinetobacter* : Touchon et al., 2014, in Archea: Wolf et al., 2016). In our *S. enterica* dataset, we evidence a clear phylogenetic signal in the gene patterns by comparing the parsimony of observed patterns to that of patterns simulated without tree (Supplementary Figure S2). Phylogenetic trees are usually supposed to represent relatedness relationships between individuals, or clonal relationship in the case of bacteria. A different view has recently emerged, suggesting that reconstructed bacterial trees reflect the structure of recombination fluxes inside the population rather than actual clonal relationships (Sakoparnig et al., 2021). While the present study does not bring new arguments supporting either of these views, we believe that the PPM model remains relevant even in the latter one. In this view, the fact that Private genes are restricted to a given clade would not be a consequence of vertical inheritance (i.e., no transfers) but of transfers restricted to a certain subpopulation, represented by a clade in the core tree. Methods exist to describe how the molecular diversity of genes is shaped by gene duplication, transfer and loss on a species tree, but these computationally intensive methods would currently not be adapted for comparative pangenomics on large datasets (Szöllosi et al., 2012; Szöllosi et al., 2015; Jacox et al., 2016; Bansal et al., 2018).

### A new pangenomic statistic to visualize goodness-of-fit

As the field of pangenome evolution is largely in its infancy, it is important to devise new statistics to visualize pangenome data and assess goodness-of-fit of models (Baumdicker & Kupczok, 2023). While the first statistics used to describe bacterial pangenomes were the core genome rarefaction and pangenome enrichment curves, the Gene Frequency Spectrum (GFS) was subsequently used to calibrate models of pangenome evolution (Baumdicker et al., 2012; Haegeman & Weitz, 2012), compare different models (Lobkovsky et al., 2013), and compare processes at work in clonal versus recombinant bacterial species (Bolotin & Hershberg, 2015).

We introduce a new statistic for representing the distribution of genes in a set of genomes: the parsimony vs. frequency plot. Each gene is represented by a point, the abscissa of which corresponds to the gene’s frequency, and the ordinate to the minimum number of gains and losses required along the species tree to generate the gene’s presence/absence pattern. A gene’s parsimony score can be seen as indicating whether the gene is preferentially present in phylogenetically close genomes or not. The parsimony vs. frequency plot is a two-dimensional statistic, and therefore contains more information than the GFS (the latter can be found by projecting the scatterplot onto the *x*-axis). It can be used to visually compare pangenomes from different populations, or to estimate the goodness-of-fit of a model, as we do.

These two statistics (GFS and parsimony vs. frequency plot), along with the visualization of some patterns with poor likelihood, help us suggest ways to improve our model. We chose to formulate a parsimonious model with a small number of gene categories to interpret the classification resulting from the model in the light of genome organization, and gene function. Some developments of the PPM model could help increase the goodness-of-fit. In particular, in our data many genes have low to moderate frequency and non-parsimonious patterns (with more than 50 gain/loss events) and many genes have medium-large frequency (carried by around 600-800 genomes), features that our model can hardly reproduce (Figures 5a and 5d). One solution would be to allow rate heterogeneity inside one or several of the three categories; this possibility is explored in section 5.2 of the Supplementary Text. However, genes with patterns that are poorly fitted often exhibit behaviours intermediate between Private and Mobile: either present at high frequency in several clades, or present at high frequency in one clade and low frequency in the rest of the tree. These could be better fitted if Private genes were allowed to be gained several times, or if genes were allowed to change categories over time: Private genes could become Persistent at some point (i.e., reduce their loss rate) or Mobile could become Private in certain clades (i.e., be lost forever when they are lost, but at a lower rate). Allowing genes to change category would also be a way to ensure stationarity in the gene number of each category; in the current version of the model Persistent genes can only decrease in number through time and Mobile genes can only increase in number.

### Gene pool model

One of the new modelling ideas introduced in this paper is the gene pool model used for Mobile genes. This model accommodates intra- and inter-species exchanges, and is both less cumbersome and more relevant than a model based on explicit exchanges between lineages (because, however large the dataset, sequenced genomes will always represent a tiny fraction of a bacterial species). Even more interestingly, our simulation study suggests that it should be possible to infer a date of introduction for each Mobile gene, if applied to a set of genomes from a recently emerged clade (see section 3 of the Supplementary Text and Figures S5 and S6 for details on arrival time inference). This would allow us to infer the temporal construction of the pangenome of the clade by progressive addition of genes. In future works, this model could be used to build a phylo-ecological model taking into account both gene dynamics and ecological niche, to study the formation of multiple “niche gene pools” as lineages move between different environments.

In conclusion, the PPM model classifies gene dynamics into three qualitatively different behaviours, infers relevant rates, and identifies hotspots of gene exchanges along the genome. Applied to the large number of bacterial whole-genome sequences, it could significantly improve our understanding of bacterial pangenome dynamics and evolution.

## Methods

### *Salmonella enterica* dataset

We applied our model to a dataset of 902 *S. enterica* genomes and one *Salmonella bongori* as outgroup, downloaded from the Refseq database. We chose to work with complete genome assemblies, as draft assemblies can cause an artificial increase of the gene number due to fragmented genes (Denton et al., 2014). We thus selected all complete *S. enterica* genomes that were available in this database, which yielded 1374 genomes (genomes downloaded the 07/09/2023, Refseq accession numbers listed in Supplementary Material). We performed the pangenome analysis using the PanACoTA pipeline (v.1.4.0, Perrin and Rocha, 2021). After quality filtering 903 genomes were kept (with a maximal distance of 0.1 between 2 genomes). Among these genomes, 624 contain one or several contigs labelled as plasmids in addition to the contig corresponding to the chromosome. 46,146 gene families were detected in this dataset. The persistent genome, that we defined as gene families having exactly one member in at least 99% of genomes, is composed of 2,887 families. The species tree was reconstructed using sequences from the persistent genome with IQ-TREE 2 (v.2.1.2, Minh et al., 2020). We used an ultrametric version of this tree obtained with LSD2 (v.2.4.4, To et al., 2016), which is shown on Supplementary Figure S1. This tree has a height of *H* = 0.03 and a total branch length of *L* = 0.84 substitutions per site.

The data that we use in order to fit our model is the reconstructed species tree, as well as the binary presence/absence matrix describing which gene is present in which genome. As the gene clustering step of the pipeline allows gene families to have more than one member per genome, the presence/absence matrix is in fact non-binary. However, our model does not account for gene duplication, thus we use the binarized version of this matrix.

### Likelihood computation

In the following, we describe how to compute the likelihood of the data according to our model. Recall that the Persistent, Private and Mobile categories are indexed by 0, 1 and 2, respectively. Let us first introduce some variables. The number of genes in the pangenome of the bacterial population described by the model is

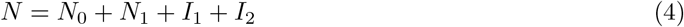

where *I*_1_ ∼ *Pois*(*i*_1_*L*) and *I*_2_ ∼ *Pois*(*i*_2_*H*). *L* is the total length of the species tree *T, H* is its height. Remember that we chose to fix *N*_1_ = *i*_1_*/l*_1_, which is the mean number of Private genes in a genome at equilibrium (Huson & Steel, 2004). The number of genes observed at the leaves is:

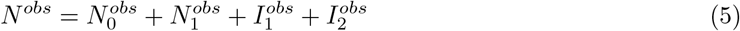

where

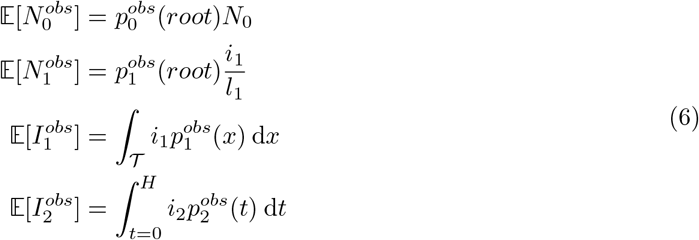

with 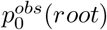 the probability for a Persistent gene to be observed a the leaves, 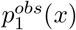 the probability for a Private gene that arrived at position *x* on the tree to be observed at the leaves, and 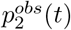 the probability for a Mobile gene that arrived in the population at time *t* to be observed at the leaves. It can be noted that the integrals used to compute 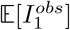 and 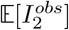 are not of the same nature. In the first one, we integrate over every possible position in the tree, while in the second one we integrate over time. In the following, we will approximate 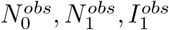 and 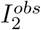 by their expectation.

Although immigrant genes from the Private category (resp. from the Mobile category) share the same realization of the random variable *I*_1_ (resp. *I*_2_), we make the simplifying assumption that all genes are independent and compute a pseudo-likelihood of observed data:

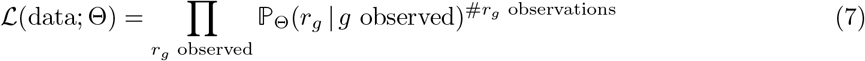

where Θ = (*N*_0_, *i*_1_, *l*_1_, *i*_2_, *l*_2_, *g*_2_, *s*_1→0_, *s*_0→1_) and *r*_*g*_ is the pattern of presence/absence of gene *g*. Then for each observed pattern the likelihood can be decomposed into three components, one for each category. As the Private category contains two types of genes (genes that were present in the ancestral genome and those that were not), we have in fact four components to compute for each patterns.

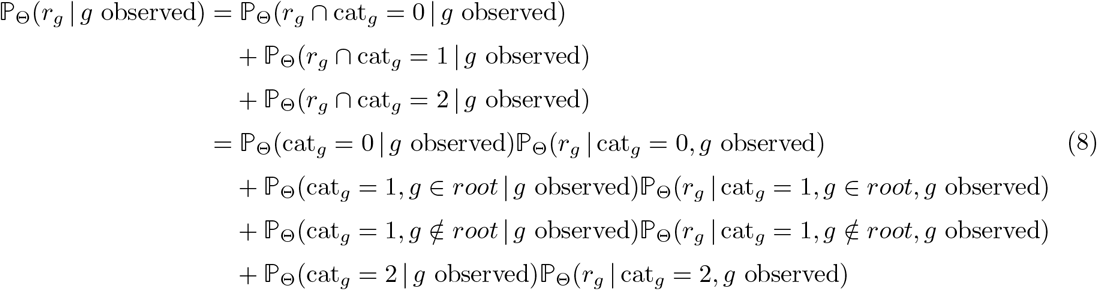

### Likelihood for Persistent genes

For a Persistent gene *g*, we denote by 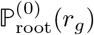 the probability to observe presence/absence pattern *r*_*g*_ given that gene *g* was present at the root (which is always the case for Persistent genes). The (0) exponent is used to indicate that this probability is computed for a Persistent gene, i.e. a gene from category 0. This quantity only depends on the loss rate *l*_0_ and the switching probability *s*_1→0_, and can be computed using Felsenstein’s pruning algorithm (Felsenstein, 1981). Formally, we have:

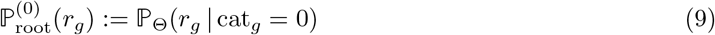

Thus:

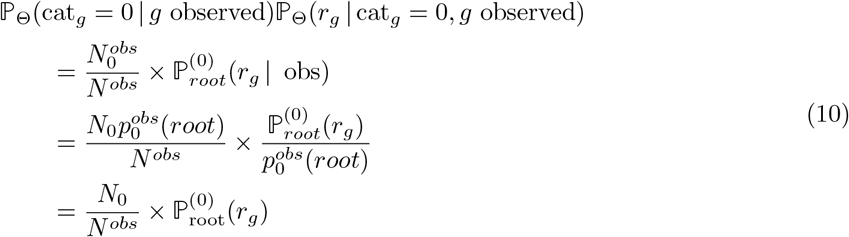

The first equality is obtained by writing the definition of each of the two factors. The second equality is obtained using equation (6) and the fact that we approximate 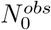 by its expectation.

### Likelihood for Private genes

For Private genes, the term corresponding to ancestral genes is similar to the one for Persistent genes:

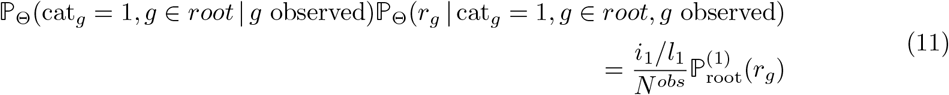

For an imported Private gene *g*, we have to integrate over all possible tree positions where this gene could have appeared. These possible positions are located on the path between the most recent common ancestor of the leaves carrying this gene (denoted by *mrca*(*r*_*g*_)) and the root of the tree. This is due to the fact that Private genes can be gained only once.

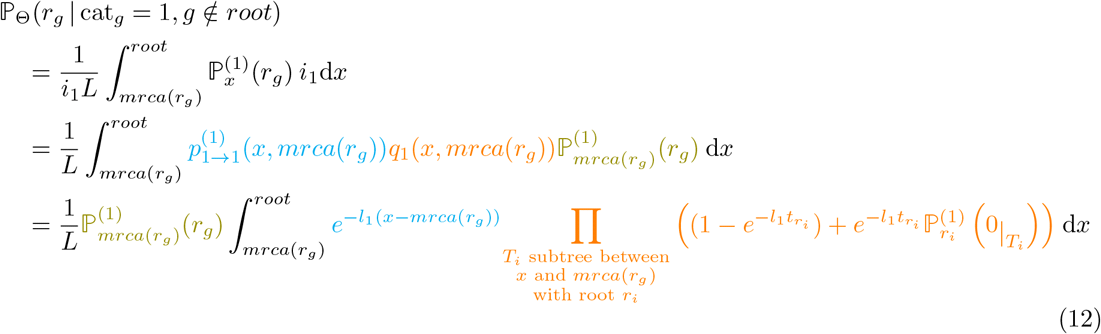

Here, the first equality integrates over all possible positions for the appearance of the focal gene on the tree, i.e. on the path between *mrca*(*r*_g_) and *root. L* is the total tree length, and 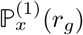 is the probability to observe pattern *r*_*g*_ knowing that gene *g* was present at position *x* in the tree.

The second equality decomposes 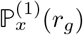 into the probability that *g* survive between its arrival at position *x* and *mrca*(*r*_*g*_) (denoted by 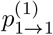 (*x, mrca*(*r*_*g*_))), times the probability that *g* was lost in all subtrees branching on the path between *x* and *mrca*(*r*_*g*_) (denoted by *q*_1_(*x, mrca*(*r*_*g*_))), times the probability to observe pattern *r*_*g*_ given that *g* was present at *mrca*(*r*_*g*_). See Figure 8a for a graphical explanation.

**Figure 8:**
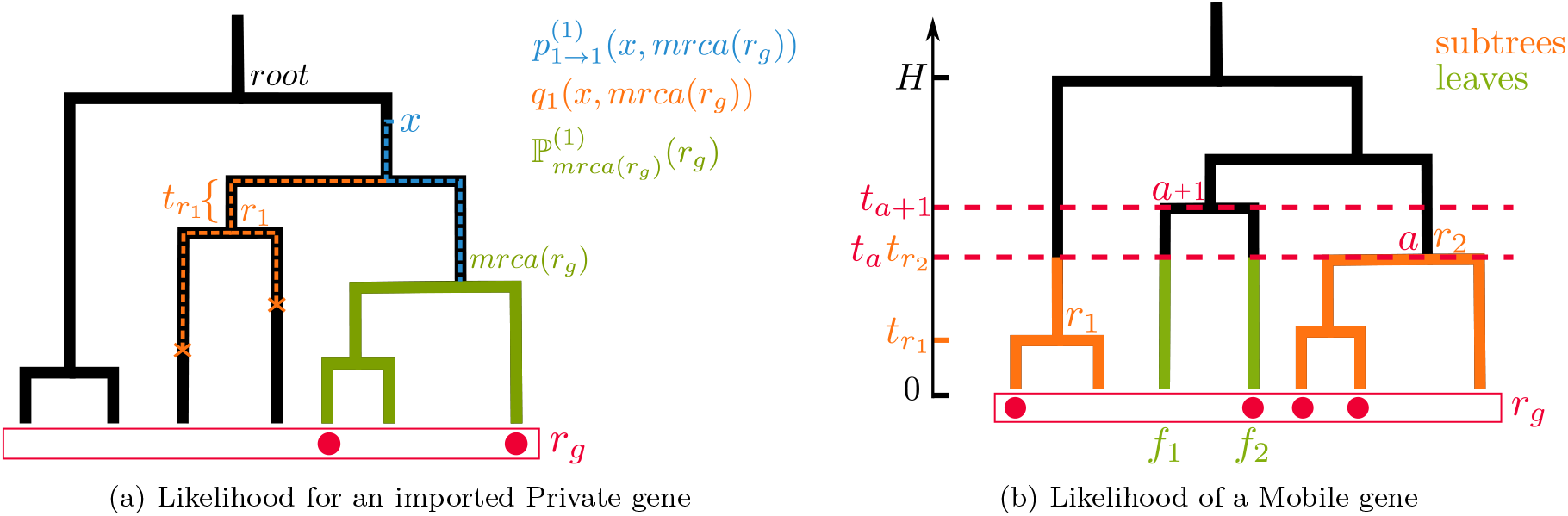
Schematic explanation of how to compute the likelihood of imported genes. (a) For imported Private genes we have to compute the probability to observe pattern *r*_*g*_ given that *g* was present at *mrca*(*r*_*g*_) (in green), times the integral over *x* of the probability to survive between *x* and *mrca*(*r*_*g*_) (in blue) times the probability to be lost in all subtrees branching between *x* and *mrca*(*r*_*g*_) (in orange). See equation (12). (b) For Mobile genes we have to sum over all time intervals of the tree (i.e., intervals between 2 nodes) the product of the probabilities to observe pattern *r*_*g*_ restricted to subtrees created by cutting the tree at this interval (in orange), times the product of probabilities to observe pattern *r*_*g*_ restricted to single leaves (in green). See equation (16).

The third equality is obtained by factoring out 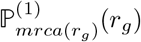 and writing the expressions for *p*_1→1_ and *q*_1_. *q*_1_(*x, mrca*(*r*_*g*_)) is the product for each subtree *T*_*i*_ branching between *x* and *mrca*(*r*_*g*_) of the probability that *g* is lost before the root *r*_*i*_ of *T*_*i*_, plus the probability that *g* survive until *r*_*i*_ and that the null pattern (denoted by 0) is observed at the leaves of *T*. 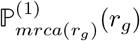 and 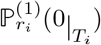 can be computed using Felsenstein’s pruning algorithm.

It follows that:

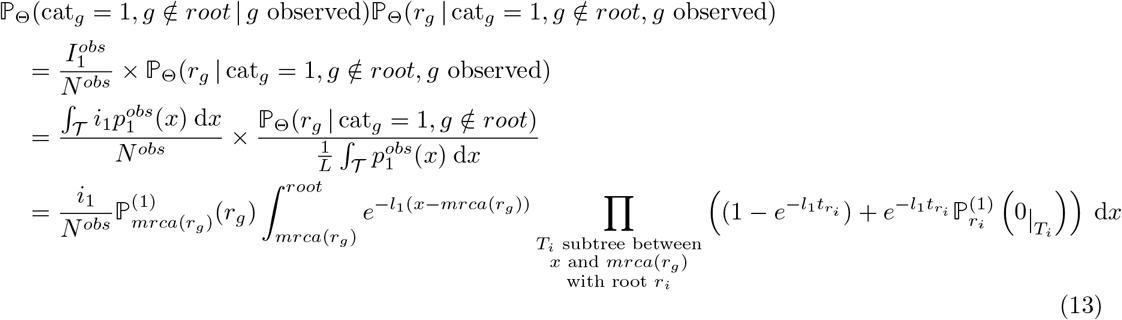

To obtain the second equality, we used the approximation of 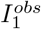 by its expectation as computed in equation (6), and the fact that 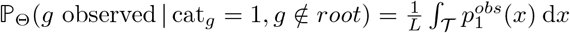.

### Likelihood for Mobile genes

For a Mobile gene, we have to integrate over the tree height for possible appearance times in the gene pool of the population. Once the gene is present in the gene pool, it evolves following a 2-state Markov model (0: absent and 1: present) with transition rates *g*_2_ and *l*_2_. The rate matrix is:

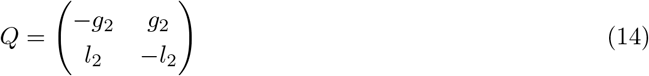

Thus the transition matrix for a time duration of *t* is:

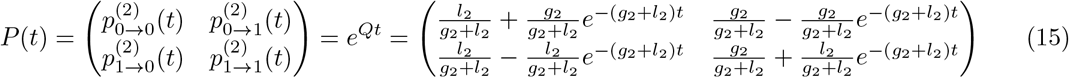

In the following time flows backward, i.e. greater time is further away in the past.

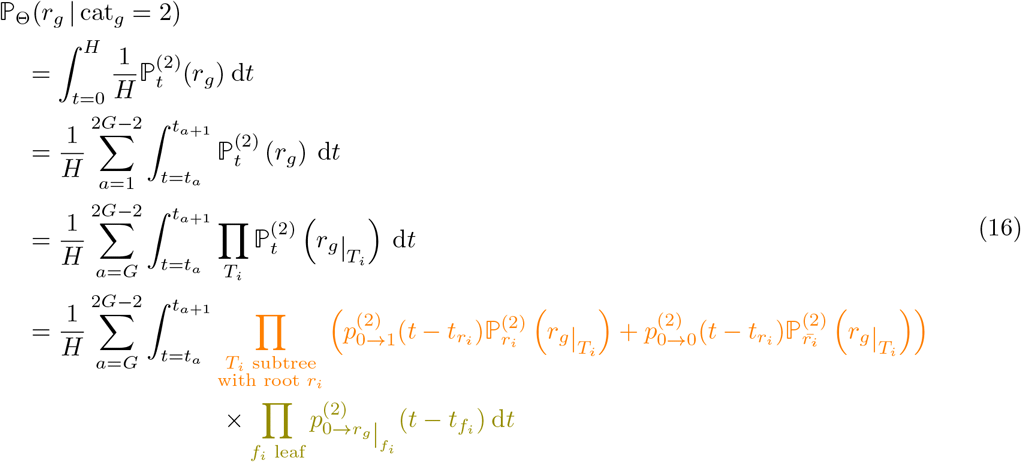

The first equality integrates over all possible heights where the gene was imported in the gene pool of the population. 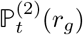 is the probability to obtain pattern *r*_*g*_ given that *g* arrived at height *t* of the tree.

The second equality decomposes the integral, summing the probability that gene *g* arrived between node *a* and node *a* + 1 over all nodes of the tree. We assume here that nodes are ordered by increasing height, i.e. *t*_1_ = 0 and *t*_2*G*−1_ = *H*. As *T* is ultrametric, we have in fact *t*_1_ = *t*_2_ = … = *t*_*G*_ = 0. In the third equality, 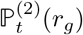 is decomposed into a product over all subtrees obtained by cutting the phylogenetic tree at time *t* (the time of appearance of the gene).

In the fourth equality, we distinguish between subtrees that are a single leaf (second product) and other subtrees (first product), see Figure 8b. In the first product, the sum represents the fact that the gene can either be gained by the focal lineage from time *t* to 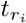 (with probability 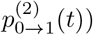, or not be gained (with probability 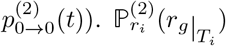 is the probability to obtain pattern *r*_*g*_ restricted to the leaves of *T*_*i*_ given that *g* was present at root *r*_*i*_ of subtree *T*_*i*_, while 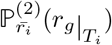 is the probability to obtain this pattern given that is was absent at root *r*_*i*_. These two quantities can be computed using Felsenstein’s pruning algorithm. 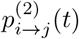 is the probability for a Mobile gene to switch from state *i* to state *j* during a time interval of length *t*, with *i, j* ∈ {0, 1}. From (15) we have:

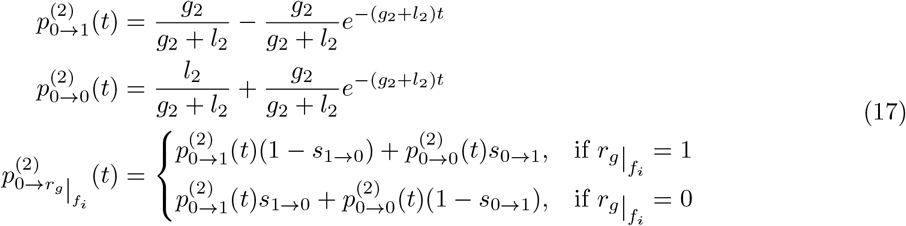

with *s*_0→1_ and *s*_0→1_ the switching probabilities. It follows that:

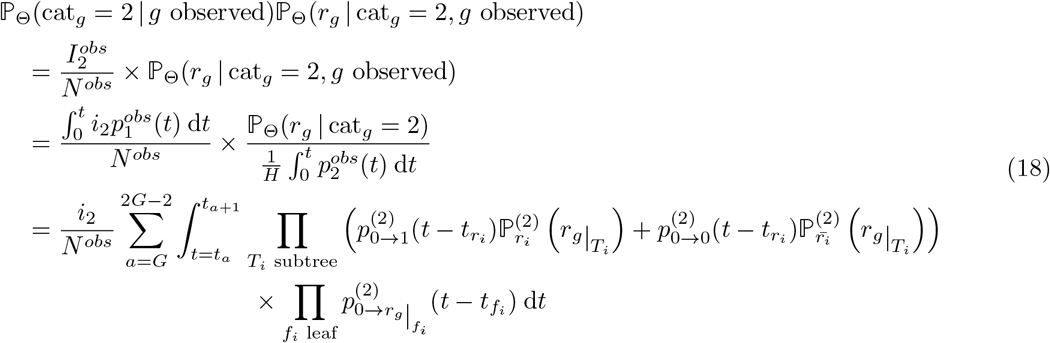

To obtain the second equality, we used the approximation of 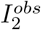 by its expectation as computed in equation (6), and the fact that 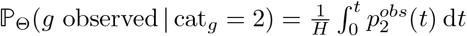.

## Inference method

### Optimization method

In order to fit our model on the *S. enterica* dataset, we computed the Maximum Likelihood parameters of the model:

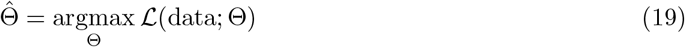

Substituting the expectations of equation (6) in equation (5), we observe that the 9 parameters (*N*_0_, *l*_0_, *i*_1_, *l*_1_, *i*_2_, *g*_2_, *l*_2_, *s*_1→0_ and *s*_0→1_) are linked by the following relation:

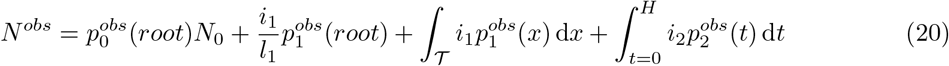

(given that 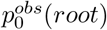 depends on *l*_0_ and 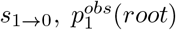 and 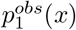 depend on *l*_1_ and *s*_1→0_, and 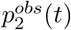 depends on *g*_2_, *l*_2_, *s*_1→0_ and *s*_0→1_). *N*^*obs*^ being the total number of observed genes in the dataset, we know its value. This relation is a constraint that we have to put on parameters to ensure that the expected number of observed genes predicted by the model coincides with the number of actually observed genes. Hence, we have only 8 degrees of freedom in the model. For example, we can express *i*_2_ as a function of other parameters:

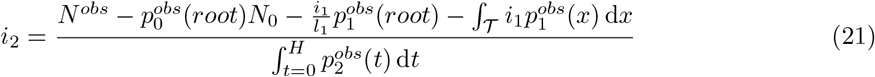

We performed likelihood maximization over the 8 free parameters using the Nelder-Mead algorithm, which is a derivative-free optimization method. As this procedure offers no guarantee to find the global maximum, we performed 10 optimizations with different, randomly chosen starting points. All 10 optimizations yielded the same result, suggesting the global maximum was reached. These optimizations took from 7 to 17 hours to run on 50 cores, depending on the starting point. The time complexity is linear in the number of gene families and quadratic in the number of genomes. The inference tool is available at https://github.com/JasmineGamblin/PPMmodelPangenome. More details on the optimization method (including the boundaries imposed on parameters and the way to draw random starting points) are given in section 1 of the Supplementary Text.

### Category assignment

Once the MLE of parameters was computed, we assigned each gene to the category with highest posterior probability, i.e. gene *g* with pattern *r*_*g*_ was assigned to category *c* such that:

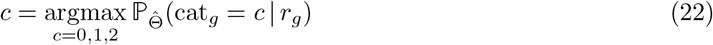

### Expected number of gains for a Mobile gene

Let *G*_*g*_ be the number of introductions for a Mobile gene *g*. By ‘introduction’, we mean a transfer between the gene pool and a lineage of the tree where this gene has never been present. We can compute *G*_*g*_ as follows, integrating over all possible arrival times in the gene pool and summing over all possible branches where an introduction could happen:

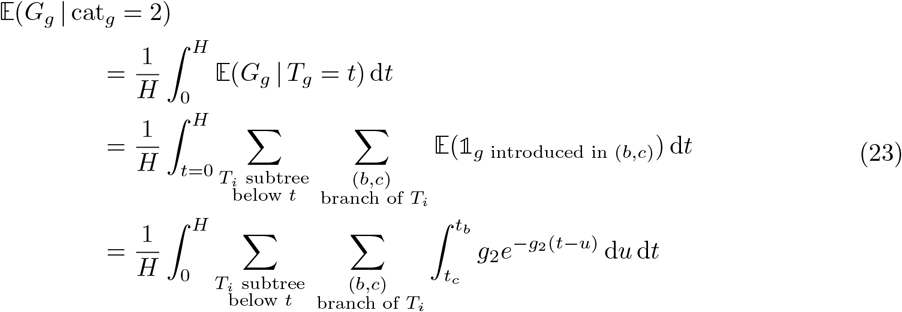

Conditionally on *g* having arrived at time *t* in the pool, the probability that it is introduced on a branch (*b, c*) below this time *t* is 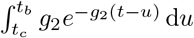. Indeed, *g*_2_ d*u* is the probability that *g* is gained during time interval d*u* and 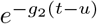 is the probability that no gain happened before time *u*.

## Acknowledgements

All authors are grateful to the Center for Interdisciplinary Research in Biology (CIRB, Collège de France) for funding. JG’s PhD scholarship was funded by the French Ministry of Research (MESRI) and by the Fondation pour la Recherche Médicale (FRM, scholarship n°FDT202304016561). The authors thank the INRAE MIGALE bioinformatics facility (MIGALE, INRAE, 2020. Migale bioinformatics Facility, doi: 10.15454/1.5572390655343293E12) for providing help and computing resources. They also thank Marie Touchon and Bastien Boussau for insightful comments about this work.

## Data availability

The inference tool for the PPM model and the code used to run simulations is available at https://github.com/JasmineGamblin/PPMmodelPangenome. Genomic data and softwares used to analyse it are referenced in the Methods section.

