## Supplementary Text for "Persistent, Private and Mobile genes: a model for gene dynamics in evolving pangenomes"

**1 Detailed inference procedure** **2**

### 1 Detailed inference procedure

The likelihood of the data is computed as explained in the Main Text. The optimization of the log-likelihood is done by the Nelder-Mead algorithm. Details of this optimization are described below.

#### 1.1 Starting point of the optimization algorithm

As explained in the Main Text, the starting point of the optimization is either randomly selected, or chosen via a pre-processing step. Random starting points were drawn from the following distributions:

- $N_0 \sim \mathcal{U}[M/3, M]$
- $l_0 \sim \mathcal{E}(L)$
- $i_1 \sim \mathcal{U}[\frac{N^{obs}}{5L}, \frac{N^{obs}}{L}]$
- $l_1 \sim \mathcal{E}(L/50)$
- $g_2 \sim \mathcal{E}(L/50)$
- $l_2 \sim \mathcal{E}(L/2500)$
- $s_{1\to 0}, s_{0\to 1} \sim \mathcal{E}(100)$

The symbol $\mathcal{U}$ denotes the uniform distribution and the symbol $\mathcal{E}$ denotes the exponential distribution. The parameters of these distributions depend on dataset characteristics, in order to automatically adapt to new datasets. More precisely, they depend on the mean number of genes per genome ($M$), the number of observed genes ($N^{obs}$) and the total branch length of the tree ($L$).

#### 1.2 Exploration of parameter space

To prevent the algorithm from exploring implausible regions of the parameter space, we placed the following constraints on the ranges of parameters, listed below:

- $N_0 \in [0, 2M]$
- $l_0 \in [0, 10/L]$
- $i_1 \in [0, 3N^{obs}/L]$
- $l_1 \in [0, 500/L]$
- $g_2 \in [0, 500/L]$
- $l_2 \in [0, 25000/L]$
- $s_{1\to 0}, s_{0\to 1} \in [0, 0.2]$

As explained in the Methods section, the estimate of parameter $i_2$ is deduced from the estimates of other parameters and $N^{obs}$, and so has to be computed at each evaluation of the log-likelihood. If the returned value for $i_2$ is negative, the function evaluation returns a very low value ($-K_2 + i_2$ where $K_2$ is taken equal to $10^9$) to drive the algorithm away from this parameter configuration. We use a threshold value of $k_1 = 10^{-4}$ on variations of the log-likelihood to terminate the optimization procedure. The values of $k_1$ and $K_2$ may have to be adapted to the size of the dataset, as bigger datasets have lower likelihood. We verified that all of the parameters inferred for the *Salmonella enterica* data lied away from these boundaries.

##### 1.3 Accelerating the likelihood computation

Computing the likelihood of our dataset given the PPM model takes about 2 minutes on 50 cores. This is due to the large number of genes in our dataset (46,146 genes, compressed into 15,192 unique presence/absence patterns), the size of our tree (902 leaves) and the fact that we have to integrate over all possible arrival times and all subtrees when computing the likelihood of Mobile genes. In order to reduce this time as much as possible, we implemented a tree data structure to store intermediate results of pruning algorithms. With this implementation, the multiple calls to the pruning algorithm requested to compute the likelihoods of Private and Mobile genes (see Main Text) never do the same computation twice during one evaluation of the dataset likelihood. Moreover, the dataset likelihood computation is easy to parallelize as the genes likelihoods are independent from each other.

##### 1.4 Avoiding underflow

During this procedure, we always compute the log-likelihood instead of the likelihood to avoid underflow (i.e., numbers that are smaller than the available precision and thus treated as zero), which are common when working with large phylogenies. Products featured in the likelihood computation are easily replaced by sums of log-likelihoods. However, sums in the likelihood require use of the so-called 'log-sum-exp trick'. Let $a = \log(x)$ and $b = \log(y)$. If we try to compute $\log(x+y) = \log(e^a + e^b)$, there is a risk of underflow if $x$ and $y$ are very small numbers. Instead, we compute $\log(x+y) = a + \log(1+e^{b-a})$. Here, we do not have to compute $x$ or $y$ directly but only their ratio $\frac{y}{x} = e^{b-a}$, which is less likely to underflow.

#### 2 Detailed simulation study

We conducted a simulation study to assess the accuracy of the inference procedure. To that end, we drew a random tree and simulated genes evolving along that tree for various sets of parameter values. Then we compared the inferred parameters to their true values on these simulated datasets (correlations between true and inferred parameters are shown in the Main Text Figure 3). The code used to generate the tree and the genes is available at https://github.com/JasmineGamblin/PPMmodelPangenome.

##### 2.1 Drawing a random phylogenetic tree

We generated a 200-leaf tree by drawing its topology from the beta-splitting trees distribution with $\beta = -1/2$ (D. Aldous, 1996; D. J. Aldous, 2001). The branching times were drawn uniformly in $[0, 1]$

and we imposed $H = 1$.

##### 2.2 Drawing the parameter values

We drew 100 sets of parameter values using the following distributions ($\log\mathcal{U}$ denotes the log-uniform distribution):

- $N_0 \sim \mathcal{U}[500, 1500]$
- $l_0 \sim \log\mathcal{U}[0.1/L, 10/L]$
- $i_1 \sim \mathcal{U}[500/L, 1500/L]$
- $l_1 \sim \log\mathcal{U}[5/L, 500/L]$
- $i_2 \sim \mathcal{U}[500/H, 1500/H]$
- $g_2 \sim \log\mathcal{U}[5/L, 500/L]$
- $l_2 \sim \log\mathcal{U}[250/L, 25000/L]$
- $s_{1\to 0}, s_{0\to 1} \sim \mathcal{E}(100)$

These distributions are designed to yield an average of a thousand genes from each category, and to span a wide range of gain and loss rates.

##### 2.3 Relative error on $g_2/l_2$ inference

We investigated what was hindering the correct inference of the ratio $g_2/l_2$ (Figure 3i in Main Text indeed shows that the inference of $g_2/l_2$ failed when its true value was smaller than $10^{-2}$). The feature of the data that was best correlated with the relative error is what we called the dispersion index: the difference between the mean frequency and the mean parsimony score, rescaled by the number of genomes. The smaller this value, the more dispersed are genes across leaves, i.e. scattered in isolated leaves rather than grouped in small clades. Figure S4 show the correlation between the dispersion index and the relative error in log scale. Given the value of the dispersion index for the *S. enterica* dataset (dashed red line), we can be confident that the ratio is well estimated in our case.

#### 3 Inference of genes' arrival time

##### 3.1 Arrival time of Private genes

As a Private gene can be gained only once along the phylogeny, it must either be already present at the root or arrive between the root and the most recent common ancestor (mrca) of genomes carrying it. If we denote by $T_g$ the time of arrival of a Private gene $g$, we can compute how long before $mrca(r_g)$ it is expected to arrive. For any points $a$ and $b$ on the ancestral lineage of $mrca(r_g)$, we denote by $q_1(a, b)$ the probability that the gene was lost in all subtrees sprouting from this lineage between $a$ and *b*. We also denote by $m$ the height of $mrca(r_g)$, and by $H$ the total tree height. Then we can express the probability density of $T_g - m$ conditional on the gene pattern $r_g$:

$$\begin{aligned}\mathbb{P}(T_g - m \in [t, t + \mathrm{d}t] \,|\, r_g) &= \frac{\mathbb{P}(r_g \,|\, T_g - m = t) \times \mathbb{P}(T_g - m \in [t, t + \mathrm{d}t])}{\mathbb{P}(r_g)} \\ &= \frac{l_1 e^{-l_1 t} q_1(m + t, m) \mathbb{P}^{(1)}_{mrca(r_g)}(r_g) \times i_1 \,\mathrm{d}t}{\int_0^\infty l_1 e^{-l_1 t} q_1(m + t, m) \mathbb{P}^{(1)}_{mrca(r_g)}(r_g) i_1 \,\mathrm{d}t} \\ &= \frac{l_1 e^{-l_1 t} q_1(m + t, m) \,\mathrm{d}t}{\int_0^\infty l_1 e^{-l_1 t} q_1(m + t, m) \,\mathrm{d}t}\end{aligned} \tag{1}$$

We will denote the denominator by $A$. Then we can express the expectation:

$$\begin{aligned}&\mathbb{E}(T_g - m \,|\, r_g, \mathrm{cat}_g = 1) \\ &\quad= \int_0^\infty t \mathbb{P}(T_g - m \in [t, t + \mathrm{d}t] \,|\, r_g) \\ &\quad= \frac{1}{A} \int_0^\infty t l_1 e^{-l_1 t} q_1(m + t, m) \,\mathrm{d}t \\ &\quad= \frac{1}{A} \left( \int_0^{H-m} t l_1 e^{-l_1 t} q_1(m + t, m) \,\mathrm{d}t + \int_{H-m}^\infty t l_1 e^{-l_1 t} q_1(root, m) \,\mathrm{d}t \right) \\ &\quad= \frac{1}{A} \left( \sum_{a=mrca(r_g)}^{root} \int_{t_a - m}^{t_{a+1} - m} t l_1 e^{-l_1 t} q_1(a, m) \,\mathrm{d}t \right. \\ &\qquad\qquad \left. + q_1(root, m) e^{-l_1 (H-m)} (H - m + \frac{1}{l_1}) \right) \\ &\quad= \frac{1}{A} \left( \sum_{a=mrca(r_g)}^{root} q_1(a, m) \left( e^{-l_1 (t_a - m)} (t_a - m + \frac{1}{l_1}) - e^{-l_1 (t_{a+1} - m)} (t_{a+1} - m + \frac{1}{l_1}) \right) \right. \\ &\qquad\qquad \left. + q_1(root, m) e^{-l_1 (H-m)} (H - m + \frac{1}{l_1}) \right)\end{aligned} \tag{2}$$

In the third equality, we write that the gene can arrive either below (left integral) or above the root (right integral). In the following steps, we decompose the left term in a sum over all nodes between the mrca and the root and integrate both the left and right terms. The quantity $A$ can be computed using the same technique:

$$\begin{aligned}A &= \int_0^\infty l_1 e^{-l_1 t} q_1(m + t, m) \,\mathrm{d}t \\ &= \sum_{a=mrca(r_g)}^{root} q_1(a, m) (e^{-l_1 (t_a - m)} - e^{-l_1 (t_{a+1} - m)}) + q_1(root, m) e^{-l_1 (H-m)}\end{aligned} \tag{3}$$

Figure S5 shows the correlation between true and inferred arrival time for Private genes simulated using the maximum likelihood estimates of parameters $i_1$, $l_1$ and $s_{1\to 0}$ from the *S. enterica* dataset. While many genes are close to the line $y = x$, some of them show an underestimated or overestimated arrival time. We believe that this can be due to specific features of the tree topology: most genes showing an underestimation (points on the bottom) arrived on the very long right branch descending from the root, while some genes showing an overestimation arrived just after the long branch which is the fifth leftmost branch from the root. There is also an over-representation of genes at very low frequency among genes with poor arrival time inference.

##### 3.2 Arrival time of Mobile genes

As we model explicitly the importation of Mobile genes into the gene pool through time, we can in theory infer the arrival dates of Mobile genes from their presence/absence pattern. Conditional on the presence/absence pattern $r_g$ of Mobile gene $g$, the distribution of its arrival time $T_g$ is computed as follows:

$$\begin{aligned}
p(T_g = t \mid r_g, \mathrm{cat}_g = 2) \\
&= \frac{1}{H} \frac{\mathbb{P}(r_g \mid T_g = t, \mathrm{cat}_g = 2)}{\mathbb{P}(r_g \mid \mathrm{cat}_g = 2)} \\
&= \frac{1}{H\mathbb{P}(r_g \mid \mathrm{cat}_g = 2)} \mathbb{P}_t^{(2)}(r_g) \\
&= \frac{1}{H\mathbb{P}(r_g \mid \mathrm{cat}_g = 2)} \prod_{\substack{T_i \text{ subtree} \\ \text{with root } r_i \\ \text{below time } t}} \left( p^{(2)}_{0\to 1}(t - t_{r_i}) \mathbb{P}^{(2)}_{r_i}\left( r_g|_{T_i} \right) + p^{(2)}_{0\to 0}(t - t_{r_i}) \mathbb{P}^{(2)}_{\bar{r}_i}\left( r_g|_{T_i} \right) \right) \\
&\qquad \times \prod_{f_i \text{ leaf}} p^{(2)}_{0\to r_g|_{f_i}}(t - t_{f_i})
\end{aligned} \tag{4}$$

$r_g|_{T_i}$ denotes the restriction of pattern $r_g$ to the leaves of subtree $T_i$, and $\mathbb{P}^{(2)}_{\bar{r}_i}(r_g|_{T_i})$ is the probability to obtain this pattern given that gene $g$ is was absent from node $r_i$. The products in the last equality run over all subtrees (including leaves) formed by cutting the tree at height $t$.

Using maximum likelihood estimates of the parameters inferred on the *S. enterica* dataset, we simulated 15,036 Mobile genes and computed the distribution of their arrival time conditional on their pattern. An arbitrary subset of the resulting distributions are shown in Figure S6a. The distributions are completely flat between height 0.0015 and 0.03, i.e. between $H/20$ and $H$. However, there are differences between genes in the time period close to the leaves, between height 0 and height $1.5 \times 10^{-4} = H/200$. In Figure S6b is represented the correlation between true arrival time (which was recorded during simulations) and inferred arrival time (computed as the mode of the conditional distribution from Equation 4). We see a somewhat satisfying correlation for genes imported very recently (between 0 and $1 \times 10^{-4}$), but earlier arrival times seem completely uncorrelated with the inferred arrival times. This is likely due to the high values of gain and loss rates for the Mobile category: the high turnover of these genes erases the signal of the introduction date very quickly, and thus the model is able to infer it approximately only for very recently acquired genes. In the light of this, the PPM model should be able to infer arrival dates of Mobile genes on datasets for which the gain and loss rates are not too high compared to tree height. In fact, we can even compute the timescale at which the gene patterns reach stationary state: according to the transition matrix (15) shown in Main Text, the characteristic time is $1/(g_2 + l_2)$, i.e. roughly $0.8 \times 10^{-4}$. This value is coherent with the above observations on simulated data.

### 4 Switching probabilities parametrisation

There is a switching term in the PPM model, that was included to account for instantaneous switching between presence and absence at the leaves of the phylogeny. As explained in the Main Text, this can be due either to sequencing and bioinformatics errors, or to rapid gene gain and loss in recent times. As we assume that Persistent and Private genes can have at most one introduction along the phylogeny, we set the instantaneous gain probability to zero for these two categories: $s^{(0)}_{0\to 1} = s^{(1)}_{0\to 1} = 0$. Four quantities remain: $s^{(0)}_{1\to0}, s^{(1)}_{1\to0}, s^{(2)}_{1\to0}$ and $s^{(2)}_{0\to1}$. We tested different parametrisation of the switching probabilities, i.e. different set of constrains on these quantities:

- **No switching**: all switching probabilities are set to zero.
- **Only bioinformatics error**: loss probabilities are equal for all categories, the gain probability is set to zero.
- **All parameters equal**: all probabilities are equal.
- **Bioinformatics error+gain**: loss probabilities are equal for all categories, the gain probability is free. This is the version presented in the Main Text.
- **Free loss probabilities**: loss probabilities are different among categories, the gain probability is equal to the loss probability of the Mobile category.
- **All parameters free**: all 4 switching parameters are free.

Table 1: Comparison of estimated parameters for several model versions, differing in the switching probabilities parametrization.

| **Model version** | **#par.** | **log-lik.** | $\hat{N}_0$ | $\hat{l}_0$ | $\hat{i}_1$ | $\hat{l}_1$ | $\hat{i}_2$ | $\hat{g}_2$ | $\hat{l}_2$ | $\hat{s}^{(0)}_{1\to0}$ | $\hat{s}^{(1)}_{1\to0}$ | $\hat{s}^{(2)}_{1\to0}$ | $\hat{s}^{(2)}_{0\to1}$ |
|---|---|---|---|---|---|---|---|---|---|---|---|---|---|
| No switching | 6 | -1,401,760 | 4,119 | 4.55 | 45,334 | 160 | 469,895 | 285 | 11,888 | 0 | 0 | 0 | 0 |
| Only bioinformatics error | 7 | -1,378,180 | 4,016 | 2.15 | 45,838 | 161 | 465,465 | 337 | 16,277 | 0.0032 | 0.0032 | 0.0032 | 0 |
| All parameters equal | 7 | -1,371,580 | 4,005 | 2.17 | 45,253 | 155 | 467,201 | 207 | 11,784 | 0.0025 | 0.0025 | 0.0025 | 0.0025 |
| Bioinformatics error + gain | 8 | -1,371,360 | 4,004 | 2.13 | 45,151 | 155 | 468,710 | 216 | 12,092 | 0.003 | 0.003 | 0.003 | 0.0022 |
| Free loss probabilities | 9 | -1,367,750 | 4,014 | 2.25 | 45,548 | 152 | 462,793 | 223 | 12,858 | 0.0021 | 0.019 | 0.0022 | 0.0022 |
| All parameters free | 10 | -1,361,600 | 4,018 | 2.27 | 45,329 | 153 | 468,244 | 180 | 8,698 | 0.0021 | 0.019 | 0.13 | 0.0022 |

Maximum Likelihood Estimates of parameters on our *S. enterica* dataset for each of these model versions are summarized in Table 1. This table shows that the other parameters of the model are not markedly affected by switching at the leaves. The estimated gain and loss rates are slightly affected as expected: when the loss probability at the leaves for a given category is smaller – e.g. when it is set to zero – the loss rate of this category is inferred to be higher. Similarly, when the gain probability is set to zero, the gain rate of the Mobile category is inferred to be higher.

Accounting for instantaneous switching at the leaves strongly increases the log-likelihood. In particular, more complex switching models (with more parameters) always show a higher log-likelihood and better AIC and BIC scores (not shown in the table). Figure S8 shows that a model with switching allows a better fit of the parsimony vs. frequency plot. The triangular shape of the bottom-right corner – where all Persistent gene are – is better reproduced than with a model without switching.

The most complex model we tested, with all free parameters, gives us insights into the relative importance of the switching processes at the leaves. The loss probabilities are ordered as loss rates across categories, with one order of magnitude between them: $s^{(0)}_{1\to0} < s^{(1)}_{1\to0} < s^{(2)}_{1\to0}$. The gain probability $s^{(2)}_{0\to1}$ is quite small, of the order of the smallest loss probability $s^{(0)}_{0\to1}$. However, the inferred parameters for this particular model need to be considered with caution. The simulation study (analogous to that on Figure 3) for this model showed that the parameter $s^{(2)}_{1\to0}$ was poorly inferred across the range of parameter values tested. This may be due to the large loss rate in this category, which generates a signal similar to that of rapid switching at the leaves.

We chose to present the version with bioinformatics error+gain in the Main Text, as this is the model with highest likelihood that also made biological sense and performed well in the simulation study. Indeed, the model with free loss probabilities but with $s^{(2)}_{0\to1} = s^{(2)}_{1\to0}$ performs well but makes little biological sense, as the gain and loss probabilities are supposed to account for different processes.

### 5 Exploration of different numbers of gene categories

We chose to build a model containing three qualitatively distinct gene categories mainly for biological reasons, as these categories represent three distinct biological behaviours that we wanted to describe. However, it could be asked whether the statistically optimal number of categories is indeed three categories for our particular dataset (the *S. enterica* dataset).
We hypothesise that genomic datasets are large and informative enough that much more complex models than three categories would be favoured by likelihood-based criterion such as AIC. The study from Zamani-Dahaj et al. (2016) found that the best model to fit their data composed of 40 cyanobacterial genomes was a model with 5 gene classes, which was also the maximum number of classes that they tested. This suggests that the optimal number of categories could be much larger for our dataset of 902 genomes. In initial explorations, we ran a model selection procedure on our dataset using IQ-TREE to find the optimal number of classes of rates of a general FMG model with rate heterogeneity (one of the models tested in Zamani-Dahaj et al. (2016)). The optimal number of classes returned by the procedure was 10, which was also the maximum number that was tested.
In the following subsections, we show that the PPM model performs better at fitting the data compared to models with only two categories, and that the Mobile genes category exhibits the highest rate heterogeneity.

#### 5.1 Statistical justification for the model with three qualitative categories

Table 2: Comparison of the likelihood and AIC of models with only two gene categories to the full model with three categories.

| | **# par.** | **log-lik.** | **AIC score** |
|---|---|---|---|
| Persistent+Private | 4 | -2,610,010 | 5,220,028 |
| Persistent + Mobile | 6 | -1,484,510 | 2,969,032 |
| Private+Mobile | 6 | -1,446,220 | 2,892,452 |
| All categories | 8 | -1,371,360 | 2,742,736 |

Log-likelihood and AIC score for models containing only two categories are shown in Table 2, and are compared with the full PPM model containing three categories. The PPM model has a lower AIC score than models with two categories, meaning that it performs better at explaining the data, even after correcting for the extra degrees of freedom.

#### 5.2 Exploration of potential rate heterogeneity within categories

We then explored the potential rate heterogeneity within each category. To this end, for each category, we selected all genes from the *S. enterica* dataset assigned to this category. We computed the maximum log-likelihood for either one or two sets of parameters on this subset. For each category, we computed the likelihood-ratio test statistic to quantify the likelihood gained from allowing rate heterogeneity (2 sets of parameters compared to one). Results are recorded in Table 3. Rate heterogeneity always considerably improves the likelihood. The largest improvement in likelihood is obtained thanks to rate heterogeneity for the Mobile gene category. This is consistent with the fact that this category contains

Table 3: Comparison of the effect of allowing rate heterogeneity in each of the three gene categories.

| | **# distinct patterns** | **log-lik. (1 set of param.)** | **log-lik. (2 sets of param.)** | $\lambda_{LR}$ |
|---|---|---|---|---|
| Persistent genes | 2,288 | -93,810 | -87,073 | 13,474 |
| Private genes | 3,302 | -376,813 | -364,389 | 24,848 |
| Mobile genes | 9,602 | -851,945 | -795,746 | 112,398 |

the highest number of distinct gene patterns. Thus, for this dataset, one could choose to fit a more complex model by allowing rate heterogeneity in priority for Mobile genes.

### 6 Simulations including duplications

In order to test if duplicated genes in the data can impact parameter inference under the PPM model, we simulated datasets containing duplications and measured how well parameters are inferred.

#### 6.1 Simulating duplications

In order to include duplications in our simulations, we added two more parameters for each category:

- $d^{(i)}$ the duplication rate for each copy of a gene from category $i$, and
- $l_c^{(i)}$ the loss rate for each copy of a duplicated gene from category $i$. Genes present in only one copy have a loss rate $l_i$.

We always chose $l_c^{(i)} \geq l_i$: it is more difficult to lose the last copy of a gene than duplicated copies. For each category, we set the ratio $d^{(i)}/l_c^{(i)}$ so that the distribution of copy numbers (pooled across all genes and all genomes) fitted the distribution observed in the *S. enterica* dataset. Precisely, to set this ratio, we fitted a geometric distribution to the numbers of gene copies; under our model, this distribution is expected to be geometric with parameter $1 - d^{(i)}/l_c^{(i)}$. The resulting ratio value is 0.0125 for Persistent genes, 0.0303 for Private genes, and 0.0161 for Mobile genes. The copy number distribution was fitted separately for each category, as we found important differences between them.

We simulated one dataset with no duplications ($l_c^{(i)} = d^{(i)} = 0$), and 4 datasets with different values for the ratio $l_c^{(i)}/l_i$, producing duplicated gene families in different proportions (recorded in Table 4).

#### 6.2 Inference results

After simulating the 5 datasets, we transformed the presence/absence matrices into binary ones and performed parameter inference under the PPM model. Table 4 shows the results of these parameter inferences, and records the error score for each dataset. The error score is computed as the mean relative error across all parameters. The proportion of duplicated families in each dataset corresponds to the proportion of gene families that are duplicated in at least one genome.

Table 4: Comparison of parameter inference on simulated data containing different proportions of duplicated gene families. True parameters used for simulations are recorded in the last line.

| $l_c^{(i)}$ **value** | **% duplicated families** | $\hat{N}_0$ | $\hat{l}_0$ | $\hat{i}_1$ | $\hat{l}_1$ | $\hat{i}_2$ | $\hat{g}_2$ | $\hat{l}_2$ | $\hat{s}_{1\to 0}$ | $\hat{s}_{0\to 1}$ | **error score** |
|---|---|---|---|---|---|---|---|---|---|---|---|
| $l_c^{(i)} = d^{(i)} = 0$ | 0 | 4,000 | 2.20 | 45,254 | 154.9 | 468,725 | 217.1 | 12,197 | 0.00304 | 0.00219 | 0.6% |
| $l_c^{(i)} = l_i$ | 5% | 4,011 | 2.07 | 45,132 | 153.7 | 468,643 | 215.1 | 11,883 | 0.00305 | 0.00219 | 0.8% |
| $l_c^{(i)} = 2l_i$ | 7% | 3,990 | 2.07 | 44,688 | 153.2 | 468,503 | 216.1 | 11,979 | 0.00299 | 0.00218 | 1.1% |
| $l_c^{(i)} = 5l_i$ | 10% | 4,001 | 2.05 | 45,218 | 152.6 | 468,976 | 215.9 | 11,935 | 0.00296 | 0.00221 | 1.1% |
| $l_c^{(i)} = 10l_i$ | 13% | 4,004 | 2.01 | 45,542 | 155.2 | 469,159 | 216.5 | 12,044 | 0.0029 | 0.00219 | 1.4% |
| true parameters | | 4,004 | 2.13 | 45,151 | 155.1 | 468,710 | 216.4 | 12,092 | 0.00304 | 0.00221 | |

The error score is always less than 1.5%, indicating that the presence of genes in multiple copies has little effect on parameter inference. Thus, for simulated datasets with copy number distributions and proportions of duplicated families reflecting those observed in the *S. enterica* dataset, gene duplications do not bias much parameter inference. As a reminder, the *S. enterica* dataset contains 8% of duplicated families, while our simulated datasets contain between 5 and 13% of such families depending on the chosen loss rate of duplicated genes.

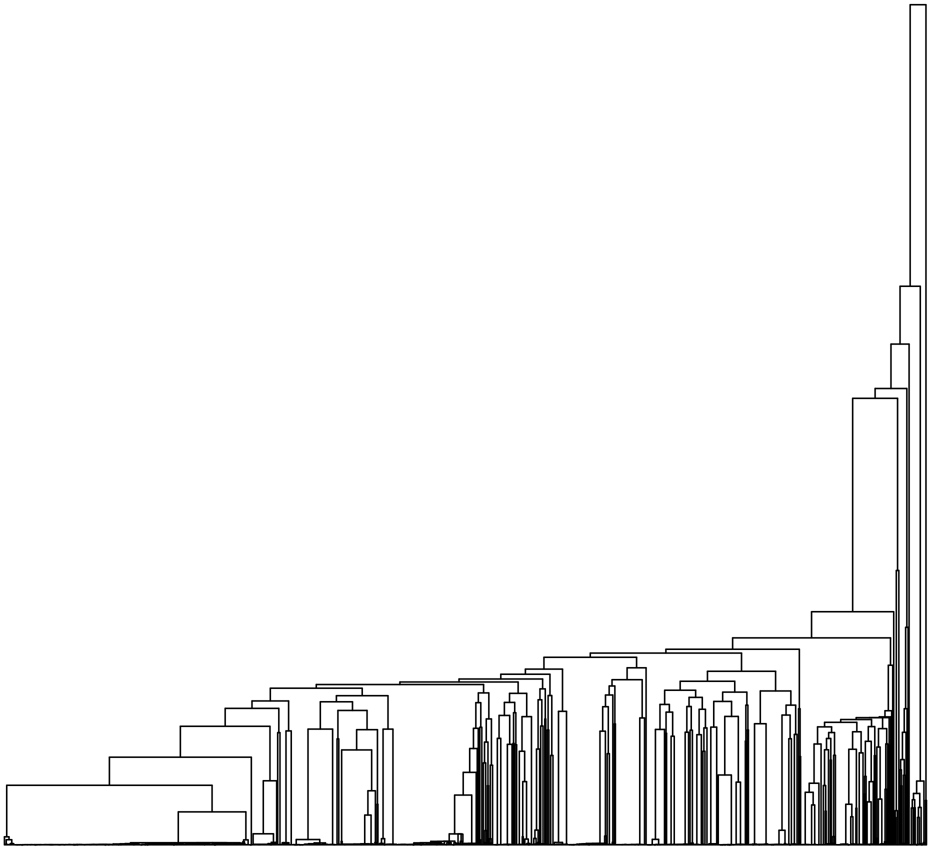

Figure S1: Phylogenetic tree of our 902 *S. enterica* genomes. It was inferred using sequences of genes present on at least 99% of genomes, with the software IQ-TREE 2. An outgroup (*Salmonella bongorii*) was used to root the tree and is not shown on the picture.

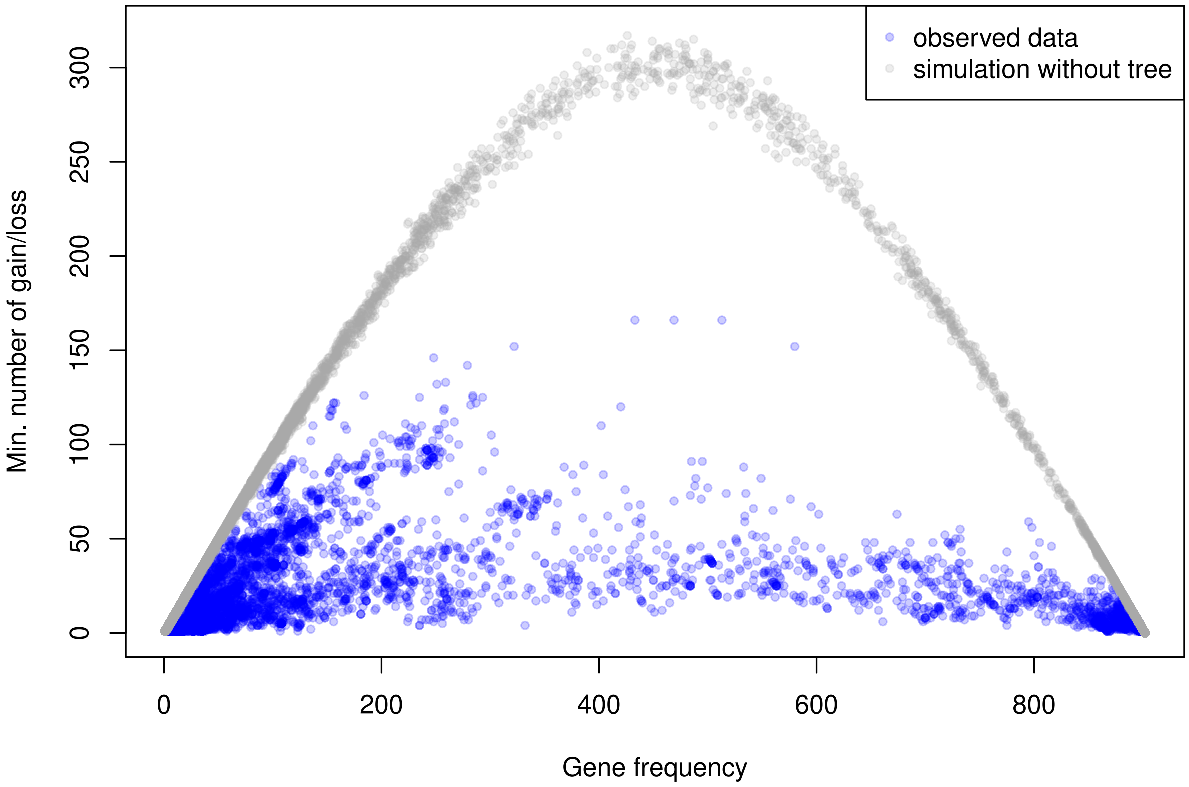

Figure S2: Blue points correspond to the parsimony vs. frequency plot of the *S. enterica* dataset. The $y$-axis is the minimum number of gain and loss events needed along the species tree to explain the presence/absence pattern of a gene with frequency given on $x$-axis. Grey points correspond to gene patterns that were randomly simulated without using the species tree (but using the same frequency distribution as in the data).

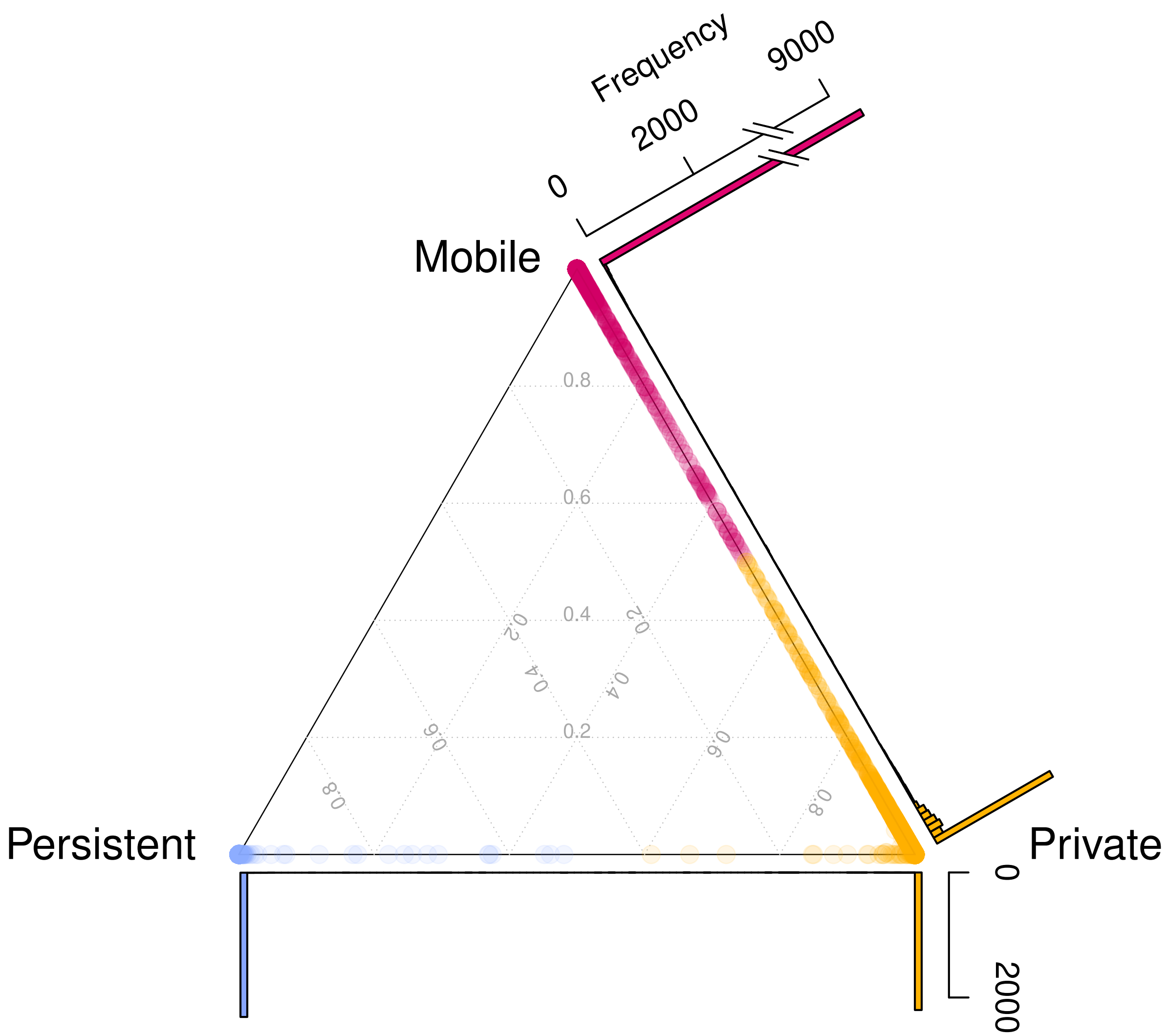

Figure S3: Ternary plot showing category assignment for genes from the *S. enterica* dataset. Each unique pattern is represented by a point, whose coordinates are the respective probabilities for this pattern to belong to each of the three categories (Persistent, Private and Mobile). Histograms show the distributions of points along two of the edges.

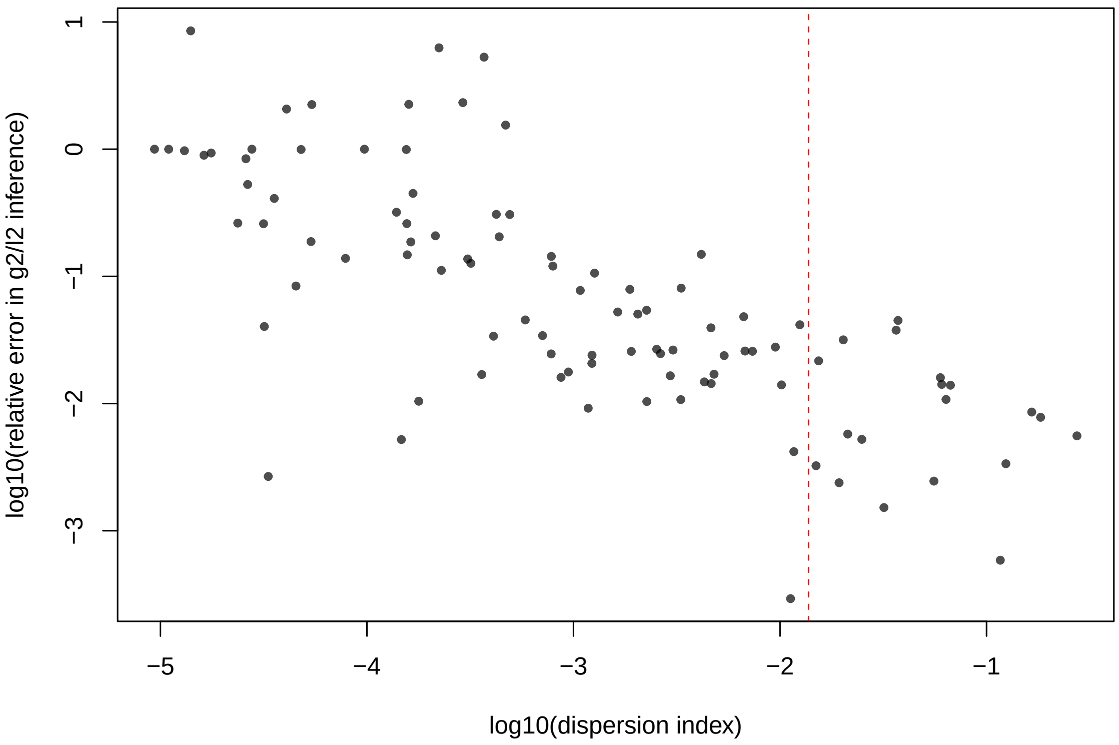

Figure S4: Relative error in the inference of $g_2/l_2$ as a function of the dispersion index, in log scale. The vertical red dashed line indicates the dispersion index of the *S. enterica* dataset.

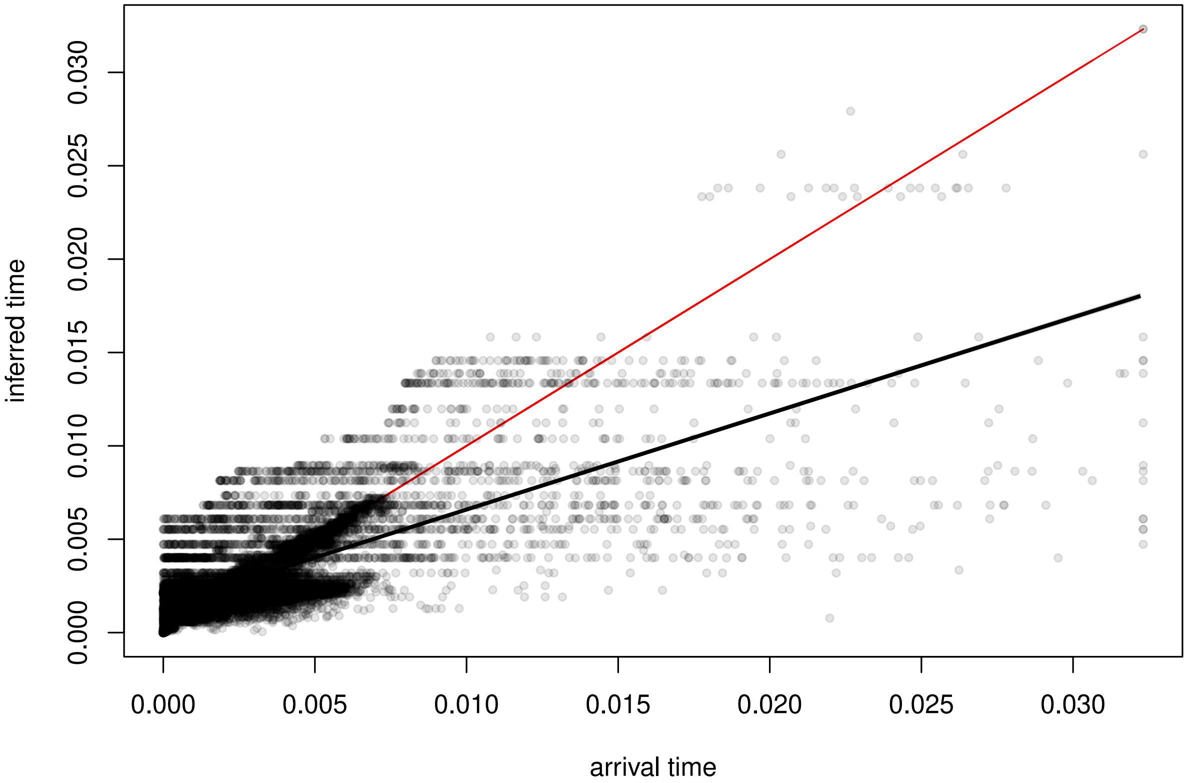

Figure S5: Correlation between true arrival time (recorded during simulations) and inferred arrival time (computed as the expected arrival time conditional on the observed presence/absence pattern) for 26,539 simulated Private genes. The line $y = x$ is plotted in red for comparison.

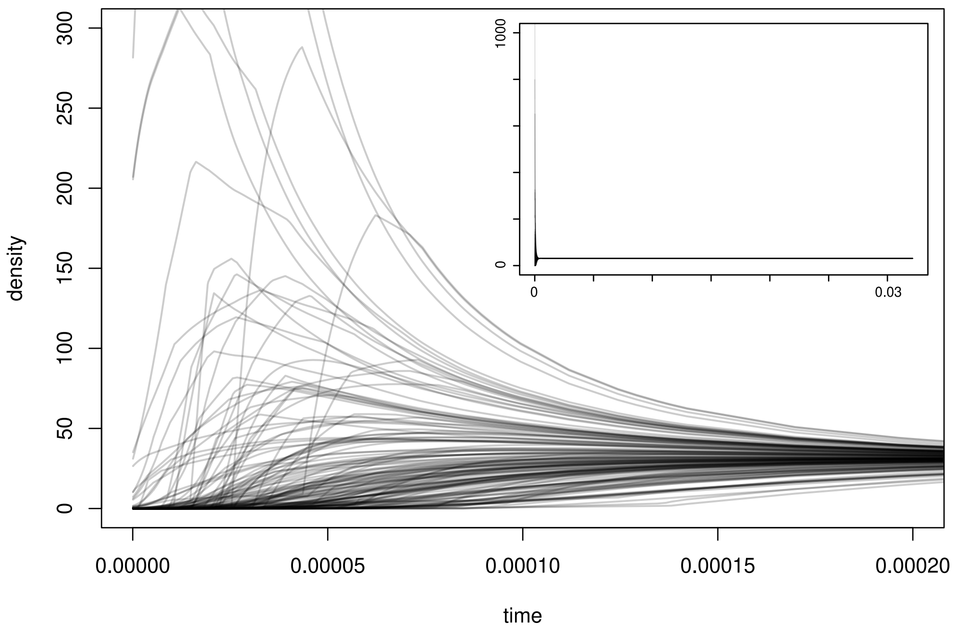

(a) Inferred distribution of arrival time

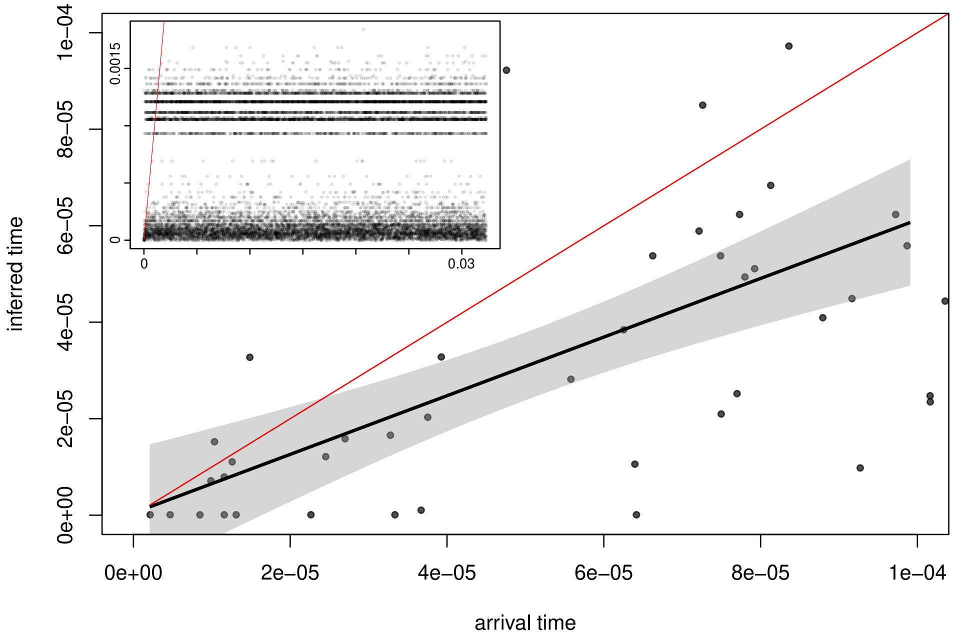

(b) Inference of arrival time

Figure S6: (a) shows the distributions of arrival time conditional on gene patterns for every $100^{th}$ gene of the 15,036 simulated Mobile genes. The main plot is an enlargement of the full distributions shown in the upper right box. (b) shows the correlation between true arrival time (recorded during simulations) and inferred arrival time (computed as the mode of the conditional distribution) for 15,036 simulated Mobile genes. The line $y = x$ is plotted in red for comparison. The main plot is an enlargement of the full picture shown in the upper left box.

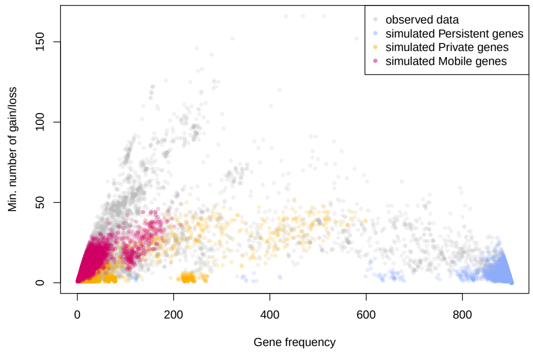

(a) Fit of the parsimony plot

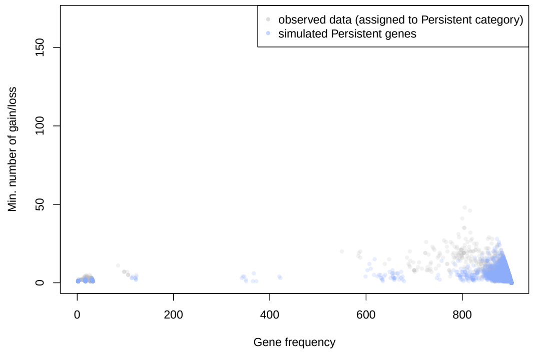

(b) Persistent genes

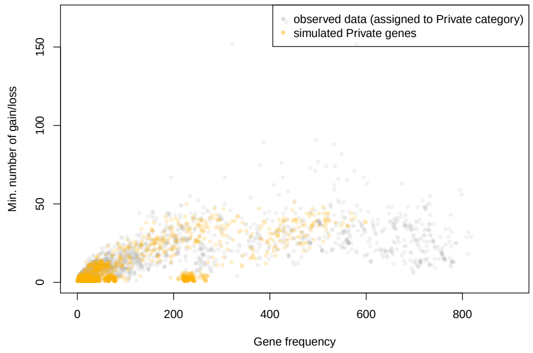

(c) Private genes

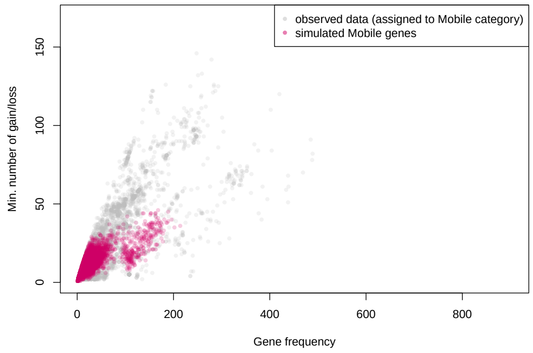

(d) Mobile genes

Figure S7: (a) Fit of the parsimony vs. frequency plot, with observed data in the background and simulated data coloured according to gene category. (b) Showing only Persistent genes (for observed data: genes assigned to this category; for simulated data: category used for simulation), (c) Showing only Private genes. (d) Showing only Mobile genes.

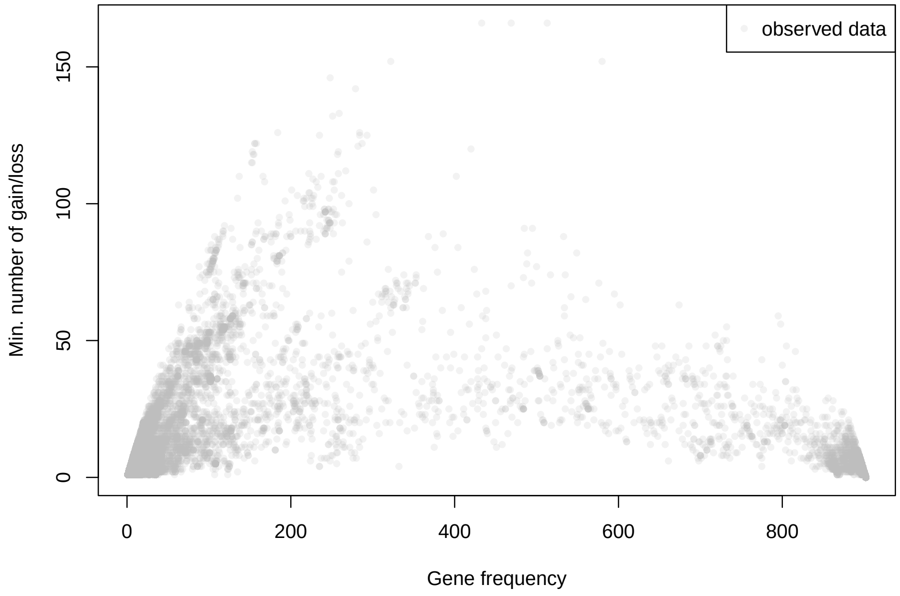

(a) Observed parsimony plot

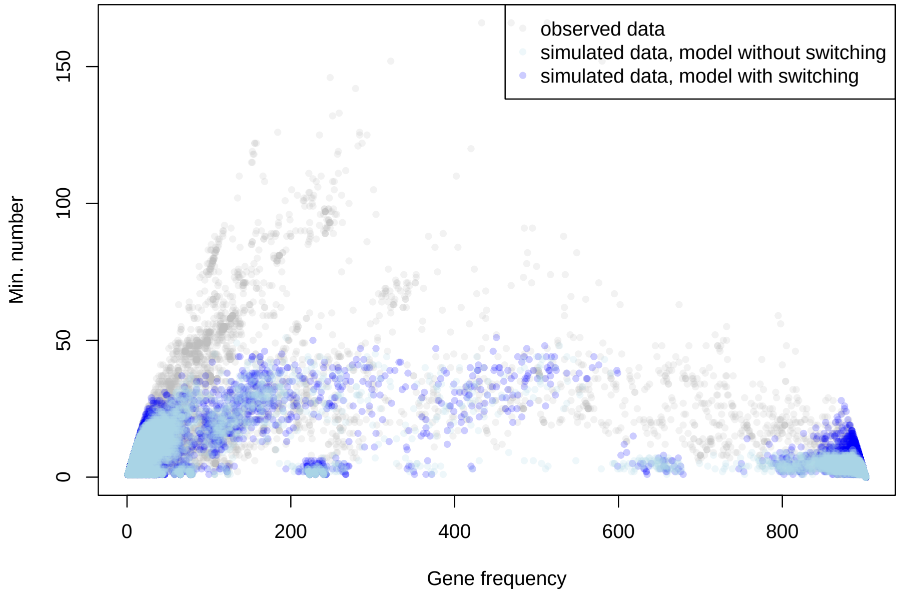

(b) Observed and simulated parsimony plots, with and without switching

Figure S8: (a) shows the observed parsimony vs. frequency plot. (b) shows the same plot, with superimposed simulated data with (blue) or without (light-blue) presence/absence switching at the leaves.

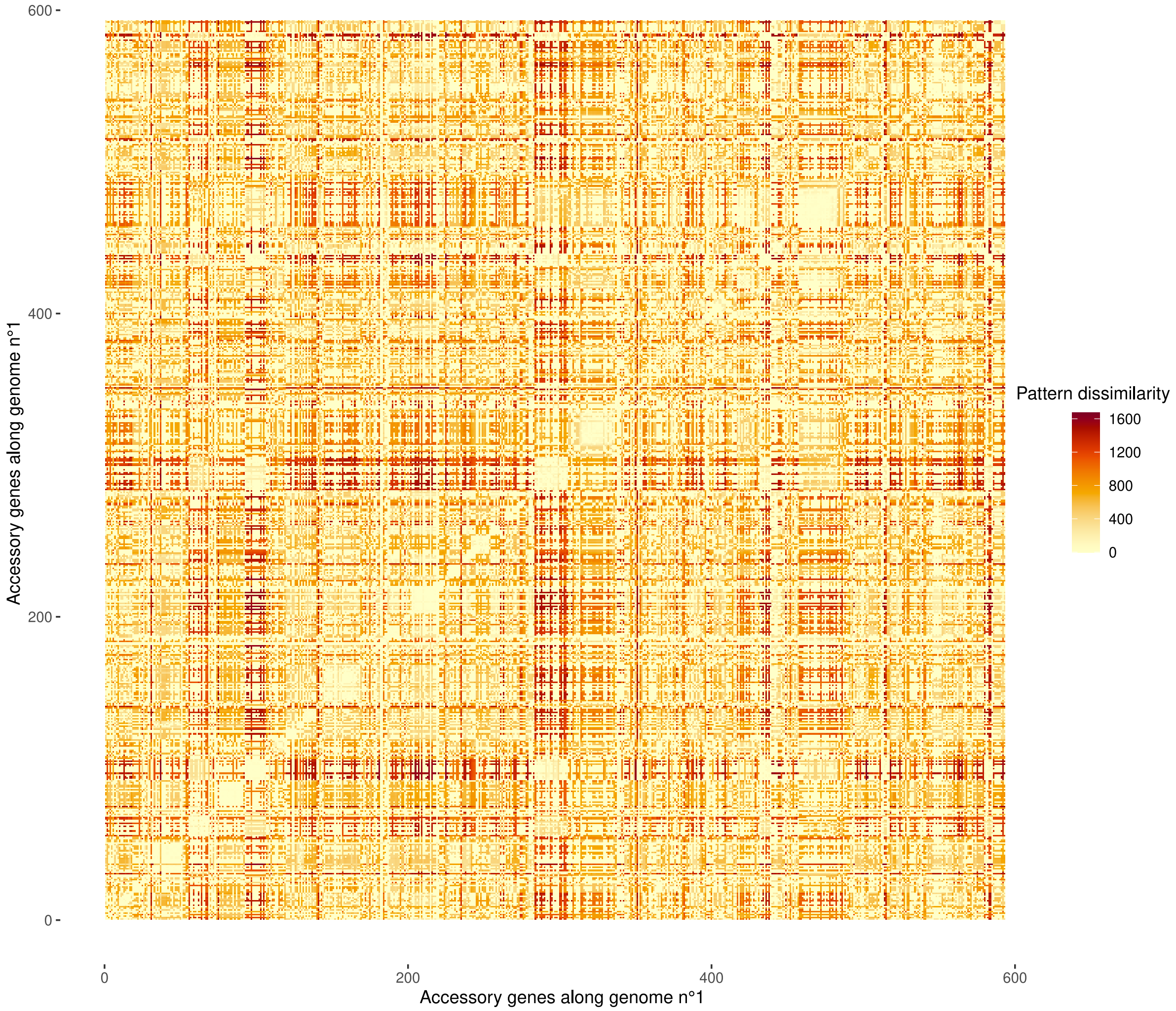

Figure S9: Heatmap showing pattern dissimilarities between accessory genes along the chromosome of genome n°1 from the *S. enterica* dataset. Pattern dissimilarity was computed as follows: for each gene, we reconstructed the ancestral states at all nodes of the species tree using maximum parsimony. The dissimilarity between two gene patterns was then calculated as the Euclidean distance between the presence/absence vectors not at the leaves, but at all nodes of the tree. This allows to correct for phylogenetic relationships among genes.
